# Nanobinders for Synaptotagmin 1 enable the analysis of synaptic vesicle dynamics in rodent and human models

**DOI:** 10.1101/2025.04.16.649111

**Authors:** Rashi Goel, Kristina Jevdokimenko, Ronja Rehm, Jannik Hentze, Paola Agüi-Gonzalez, Momchil Ninov, Felix Lange, Agata Witkowska, Svenja Bolz, Francesca Pennacchietti, Martina Damenti, Natalie Kaempf, Vladimir Khayenko, Carles Calatayud, Viveka Nand Malviya, Natali L. Chanaday, Emma Scaletti Hutchinson, Hao Liu, Kirsten Weyand, Daniela Ivanova, Tristan P. Wallis, Christopher Small, Hans M. Maric, Merja Joensuu, Michael A. Cousin, Frédéric A. Meunier, Patrik Verstreken, Ilaria Testa, Ege T Kavalali, Volker Haucke, Stefan Jakobs, Henning Urlaub, Nils Brose, Benjamin H. Cooper, Pål Stenmark, Felipe Opazo, Reinhard Jahn, Silvio O. Rizzoli, Eugenio F. Fornasiero

## Abstract

Synaptic neurotransmission is a critical hallmark of brain activity and one of the first processes to be affected in neural diseases. Monitoring this process, and in particular synaptic vesicle recycling, in living cells has been instrumental in unraveling mechanisms responsible for neurotransmitter release. However, currently available reporters suffer from major limitations such large probe size or lack of suitability for human neurons, hampering the understanding of human synaptic pathophysiology. Here we describe the NbLumSyt1 toolkit, a panel of nanobody-based affinity probes targeting the luminal domain of the synaptic vesicle protein Synaptotagmin 1 (Syt1). These new tools enable quantitative, non-invasive imaging and functional interrogation of synaptic transmission in human neurons, with unprecedented precision, versatility and cost efficiency, in technologies ranging from fixed-and live-cell super-resolution imaging to electron microscopy and mass spectrometry. Overall, NbLumSyt1 nanobinders provide a valuable platform for human synaptic physiology and pathophysiology, benefiting fundamental neuroscience and translational efforts to study and develop treatments for brain-related disorders.

## Introduction

Synaptic vesicle (SV) recycling, one of the most complex and well-regulated processes at neuronal presynapses, is responsible for the efficient and rapid release of neurotransmitters by exocytosis and the subsequent reuse of SVs ^1^. It is essential for brain function, ensures the maintenance of neuronal homeostasis, and it plays a leading role in cognitive processes such as learning and memory and in keeping neuronal circuits functionally intact ^2^. Many neurodevelopmental and neurodegenerative diseases are linked to protein mutations essential for SV recycling ^3,4^. Rodent models have contributed significantly to our understanding of synaptic physiology, but growing evidence points to distinctive features of human neurons that are not fully captured in rodents ^5^, making direct investigation in human cells essential for studying synaptic pathologies linked to cognitive decline. Despite its significance, little is known about SV recycling in human neurons, partly because appropriate tools are lacking for analyzing this process.

Synaptotagmin 1 (Syt1) is a calcium-binding protein that plays a critical role in both exocytosis and endocytosis of SVs ^6^, and that has been used as key target for studying SV recycling ^7^. Although, excellent antibodies (IgGs) are available for probing the extracellular domain of Syt1 and studying SV recycling in rodents ^8^, they only show limited specificity in human neurons. Moreover, IgGs are limited by their relatively large size (linkage-error) and bivalency. For instance, targeting multiple IgGs (diameter ∼9-14 nm) to the interior diameter of SVs (∼40 nm ^9–12^) is challenging. In addition, this can artificially cluster unfixed molecules in live assays ^13^.

Camelid single-domain antibodies, often also referred to as nanobodies, offer a promising alternative to conventional IgGs. With their small size of ∼15 kDa, nanobodies can reach epitopes sterically difficult to access with conventional antibodies ^14^. Moreover, excellent stability, ease of engineering, and low-cost production render nanobodies attractive probes for both basic research and clinical applications ^15,16^.

In this work, we describe the generation and validation of a panel of nanobodies selective for the luminal domain of Syt1, which we optimized into a diverse array of probes collectively referred to as the NbLumSyt1 toolkit. These nanobodies allow the precise and minimally invasive study of SV recycling and neuronal function in both rodent and human neurons. Our NbLumSyt1-based probes display excellent performance in live-cell labeling and can be adapted to various experimental workflows including single-molecule tracking, super-resolution imaging modalities and live-cell proteomic mapping, circumventing classical limitations of conventional IgGs.

## Results

### Development of a nanobody targeting the luminal domain of Syt1

A nanobody library was generated from peripheral blood mononuclear cells from alpacas immunized with a recombinant protein fragment corresponding to the luminal domain of rat Syt1 (**Fig. 1a**). Clones were screened for endocytic uptake in cultured hippocampal neurons (**Supplementary Fig. 1a**). Clone 1F12 displayed a clear synaptic localization and was named NbLumSyt1. A version of NbLumSyt1 fused to a HALO-Tag ^17^, allowed live-cell imaging with high signal-to-noise ratios (**Fig. 1a, b**). The NbLumSyt1-HALO specifically labeled presynaptic boutons, colocalizing with the presynaptic scaffold Bassoon and, labeling both excitatory (VGLUT-positive) and inhibitory (VGAT-positive) synapses (**Fig. 1c**). In Western Blot (WB) experiments, the sensitivity of the nanobody towards purified Syt1 was comparable to that of an established monoclonal antibody raised against the same antigen (**Fig. 1d**). No specific labeling by NbLumSyt1, fused to a GFP-tag, was observed after incubating neurons from Syt1 knockout (KO) mice ^18^, confirming specificity as also further confirmed by immunoprecipitation experiments, in which endogenous Syt1 was enriched from wildtype (WT) but not from Syt1-KO brain extracts (**Fig. 1e, f**).

**Figure 1:**
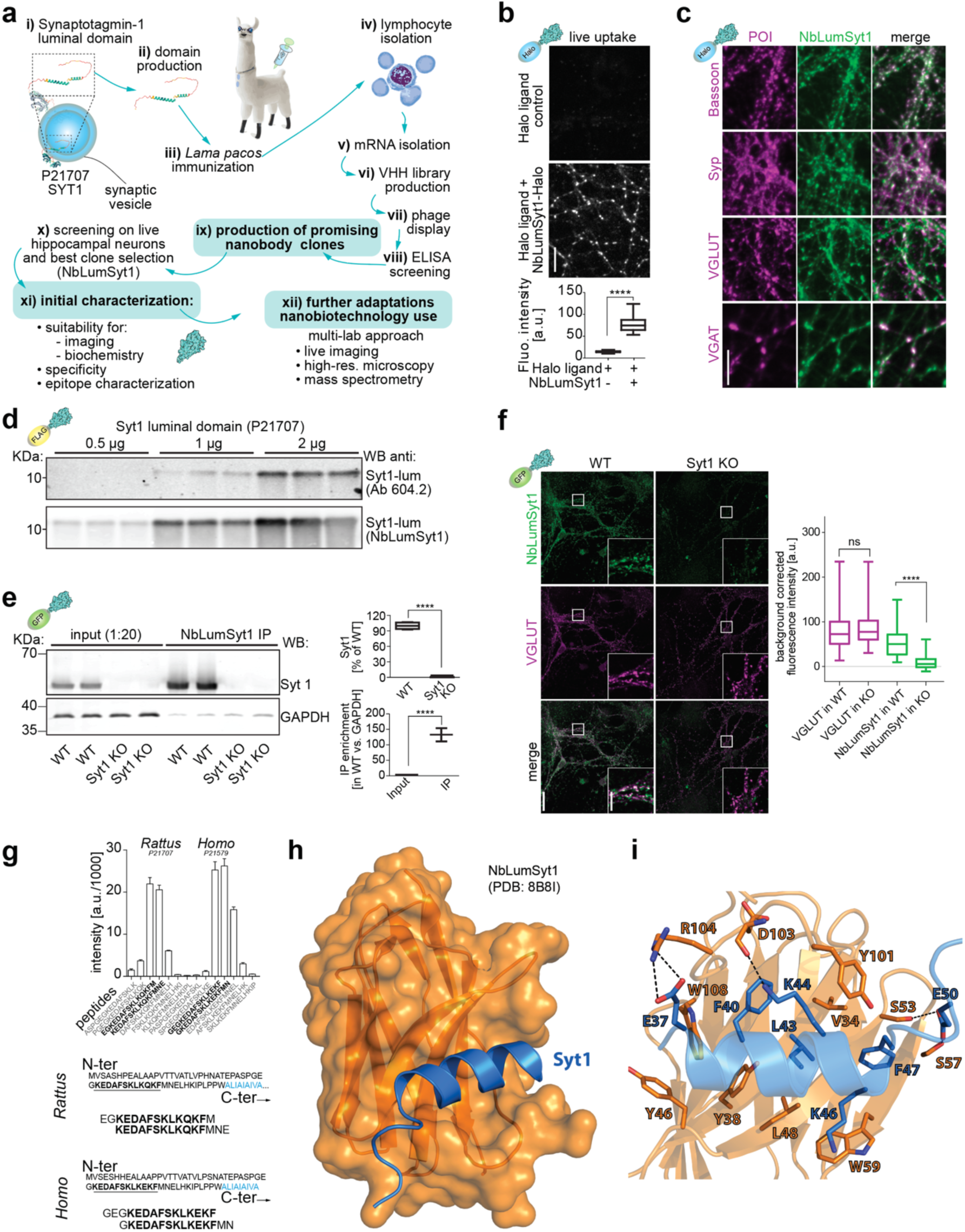
Development and characterization of a nanobody against the luminal domain of the calcium sensor Synaptotagmin 1. **a**) Schematic representation of nanobody selection and characterization. **b**) Immunofluorescence images of hippocampal neurons upon live labelling with the NbLumSyt1 fused to the Halo-Tag. This tool can be used to label actively recycling synaptic vesicles and provides excellent signal-to-noise images. **c**) Upon live imaging and retrospective immunofluorescence, the NbLumSyt1 colocalizes with the presynaptic scaffold protein Bassoon, and labels synaptic boutons, including excitatory (vGlut^+^) and inhibitory (vGAT^+^) boutons. **d**) Western blot analysis of purified Syt1. The nanobody can be used in WB applications and recognizes increasing concentrations of purified Syt1. **e-f**) Immunoprecipitation of Syt1 from WT or Syt1 KO mouse primary neurons. The nanobody is specific as it shows virtually no signal in Syt1 KO neurons in fluorescence imaging, allows to immuno-precipitate (IP) and enrich the endogenous Syt1 and shows no enrichment in Syt1 KO neurons. **g**) A peptide microarray binding assay, with synthesized and immobilized rat and human Syt1 protein sequences, identified the region of Syt1 that the nanobody binds to in both species. **h**) The crystal structure of the nanobody (orange) bound to Syt1 (blue). **i**) Specific amino acids involved in nanobody-Syt1 binding (PDB: 8B8I). Scale bars: 10 µm in b, c and f-inset; 30 µm in f.

Epitope mapping of NbLumSyt1 with a peptide microarray ^19^ identified a luminal region of Syt1 conserved between rodents and humans (**Fig. 1g**). Note that immunoreactivity was lost after chemical fixation possibly attributed to derivatization of conserved lysine residues in the epitope (**Supplementary Fig. 1b, c**). For further characterization, we solved the structure of the NbLumSyt1 in complex with the identified Syt1-peptide using X-ray crystallography at a resolution of 2.75 Å (**Fig. 1h, i**; PDB: 8B8I). The NbLumSyt1 displays a canonical IgG fold, where a disulfide bond between Cys23 and Cys96 connects two β-sheets. The Syt1 peptide forms a helical structure and binds to a shallow pocket on the surface of the nanobody. This area is primarily comprised of complementarity-determining regions (CDRs) 1, 2 and 3. The Syt1 peptide is positioned by a salt bridge between Glu37 and Arg104 and hydrogen bond interactions between Lys44 and Asp103 and Glu50 with Ser53/Ser57 of NbLumSyt1. Residues Phe40, Leu43 and Phe47 from Syt1 form hydrophobic interactions with several hydrophobic amino acids in the nanobody, including Val34, Tyr46, Trp59, and Trp108. Analysis of the nanobody binding interface with protein interfaces, surfaces, and assemblies (PISA ^20^) demonstrates a solvation-free energy gain of 10.3 kcal/mol, with an interface area of 772 Å^2^.

To further showcase the use of nanotools for studying SV recycling dynamics, we used the NbLumSyt1-HALO to examine the spatial distribution of SVs under various experimental conditions and revisited previous findings on SV intermixing ^21^. Labeling with NbLumSyt1-HALO provided clear visualization of SV populations relative to Bassoon. Pre-alignment and averaging analyses showed no significant changes in the distribution of proximal (<300 nm) or distal (400–800 nm) SV pools regardless of whether the neurons were stimulated for 2 seconds or 30 second, suggesting rapid mixing during recycling (**Supplementary Fig. 2a-f**). When assessing the localization and synaptic targeting of SVs of different molecular ages ^22^, younger SVs (1-hour after recycling) exhibited a more localized distribution near the active zone, whereas older SVs (18-hour after recycling) displayed a broader distribution, suggesting distal re-localization of aged vesicles after several hours (**Supplementary Fig. 2g-j**).

Overall, these data demonstrate that NbLumSyt1-based tools are suitable for investigating SV recycling dynamics, with wide ranging applications.

### 3D ultrastructural analysis of synaptic vesicle recycling

For electron microscopy applications, we generated a gold-labeled version of NbLumSyt1 by fusing it to ALFA-tag ^23^ and allowing complexation with a NbALFA-Gold. Cultured hippocampal neurons were incubated with the labeled complex and then analyzed by EM tomography. The high spatial resolution of internalized gold-label revealed their localization in vesicles within presynaptic terminals (**Fig. 2**). Compared to controls treated with a nonspecific NbALFA-Gold, a significantly higher percentage of tomograms showed intravesicular gold-labeling with the NbLumSyt1-ALFA-tag:NbALFA-Gold complex, along with an increased ratio of intracellular to extracellular gold-labeling (**Fig. 2a-f**). Absolute counts of intracellular and extracellular gold particles further confirmed these results. Gold particles were present inside the majority of clear core SVs (CCSVs) but also found in endosomes and multivesicular bodies (**Fig. 2g**). At high magnification, individual SVs containing luminal gold particles were resolved, and tomographic reconstructions showed their accumulation near the active zone (**Figures 2h-n**). These gold-labeled vesicles, hereafter referred to as CCSV^G^s, were enriched near active zone (AZ) release sites compared to unlabeled SVs, consistent with their preferential recruitment during recycling (**Fig. 2o-p**). Additional tomograms provided further evidence for the specificity of gold-labeling. Individual CCSV^G^s containing single or clustered gold particles were quantified, and unbiased volume estimates demonstrated a linear relationship between the number of luminal gold particles and estimated gold cluster volume (**Supplementary Fig. 3a-c**). Orthogonal views of single luminal gold particles further validated their resolution within individual vesicles and demonstrating that ET can resolve and quantify single luminal gold (3 nm; **Supplementary Fig. 3d-f**).

**Figure 2:**
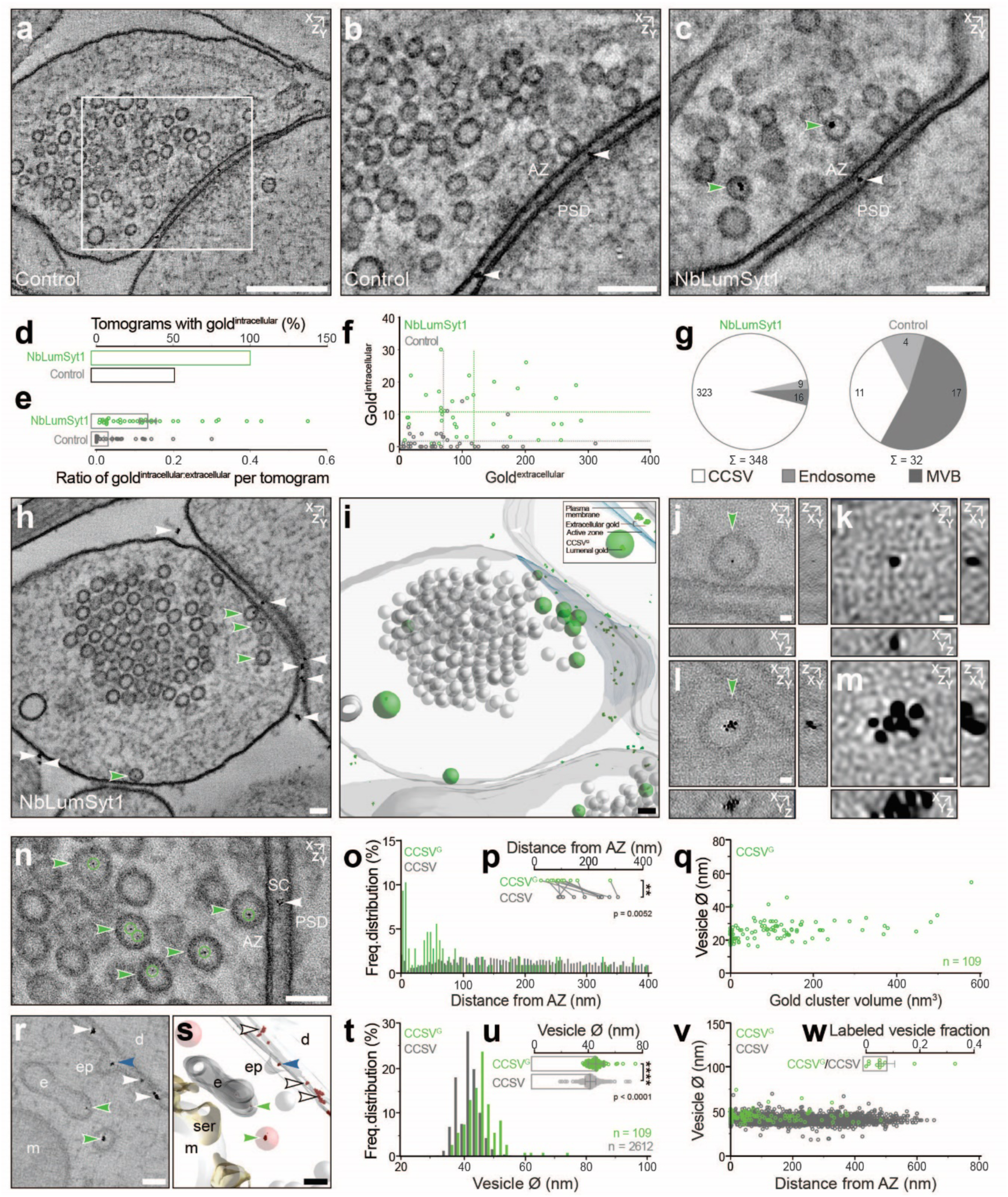
3D Ultrastructural analysis of live-cell uptake of the NbLumSyt1:NbALFA-Gold complex. **a-c**) Electron tomographic sub-volumes acquired from synapses at 36 000 x magnification (binned voxel size x,y,z = 1.2 nm; projection through 5 tomographic slices) from ALFA-Gold (a, b; Control) and NbLumSyt1:NbALFA-Gold complex (c; NbLumSyt1) treated conditions. The white box in a represents the active zone release site enlarged in b. Note that the plane chosen in b differs from panel a. **d**) Percentage of tomograms with intracellular gold (NbLumSyt1, 100%; Control, 53,85%). **e**) Ratio of intracellular gold (Gold^intracellular^) to extracellular gold (Gold^extracellular^) for each tomogram. **f**) Absolute numbers of gold clusters found intracellularly versus extracellularly for each tomogram. Dotted lines indicate mean values for control (grey; Gold^intracellular^ mean ± SEM = 1.95 ± 0.52; Gold^extracellular^, mean ± SEM = 70.60 ± 10.40) and NbLumSyt1:NbALFA-Gold complex (green; Gold^intracellular^ mean ± SEM = 11.00 ± 1.24; Gold^extracellular^, mean ± SEM = 118.10 ± 14.07) conditions. **g**) Presynaptic distribution of gold clusters within different organelles: Clear core SVs (CCSVs, white), endosomes (light grey) and multivesicular bodies (MVB, dark grey). **h-k**) Electron tomographic sub-volume acquired from a synapse at 57 000 x magnification (binned voxel size x,y,z = 1 nm; projection through 20 tomographic slices) from the NbLumSyt1:NbALFA-Gold complex (h; NbLumSyt1) treated condition and corresponding 3D model (**i**). Clear-core vesicles, CCSV (grey spheres); Gold-labeled clear-core vesicles, CCSV^G^ (green spheres); gold clusters (green); active zone, AZ (blue); plasma membrane (grey); white and green arrowheads indicate extracellularly and intracellularly localized gold clusters, respectively]. (**j-m**) Examples illustrating a CCSV^G^ containing either a single (j) or multiple (l) 3 nm gold particles as visualized in orthogonal X,Y,Z orientations (k, m, Enlargements; unbinned voxel size x,y,z = 0.5 nm; volume represents projection through 6 tomographic slices). **n)** Tomographic sub-volume (binned voxel size x,y,z = 1 nm; volume represents projection through 20 tomographic slices) revealing the tendency for gold-labeled vesicles to accumulate in close proximity to active zone release sites. CCSV^G^, green arrowheads; luminal gold particles, green circles; extracellular gold, white arrowheads. **o,p**) Spatial distribution of gold-labeled clear-core vesicles (CCSV^G^, green) and unlabeled vesicles (CCSV, grey) within 0 to 400 nm from the AZ (5nm bins) (**o**). Mean distance of CCSV and CCSV^G^ to the AZ for each tomogram (p) (CCSV^G^, mean ± SEM = 106.50 ± 20.72; CCSV, mean ± SEM = 199.30 ± 23.53; Paired Student’s t-test, p = 0.0052). **q**) Scatter plot illustrating relationship between CCSV^G^ diameter (Ø) and luminal gold cluster volume. **r-s**) Tomographic sub-volume (r; binned voxel size x,y,z = 1.2 nm; volume represents projection through 6 tomographic slices) and 3D model (s) in which multiple gold-labeled endocytic intermediates are captured (white arrowheads, extracellular gold; green arrowheads, endocytosed gold; blue arrowhead, gold localized to developing lumen of an endocytic pit). **t, u**) Frequency distribution (t, 2 nm bins) and corresponding scatterplot (u) indicating all quantified CCSV^G^ (green, n=109 vesicles) and CCSV (grey, n=2612 vesicles) diameters (CCSV^G^, mean ± SEM = 45.71 ± 0.56; CCSV, mean ± SEM = 41.75 ± 0.08; Kolmogorov-Smirnov test, p < 0.0001). **v**) Spatial distribution of CCSV^G^ (green) and CCSV (grey) relative to the AZ and their respective vesicle diameters (Ø). **w**) Relative proportion of CCSV^G^ to CCSV for each tomogram. Abbreviations: Active zone, AZ; synaptic cleft, SC; postsynaptic density, PSD; dendrite, d; endocytic pit, ep; endosome, e; smooth endoplasmic reticulum, ser; mitochondria, m. Scale bars: **a**, 250 nm; **b**, **c**, 100 nm; **h**, **i**, **n**, **r**, **s**, 50 nm; **j**, **l**, 10 nm; **k**, **m**, 3 nm. Error bars indicate mean ± SEM; ∗p < 0.05; ∗∗p < 0.01; ∗∗∗∗p < 0.0001. (**a-g**) 36 000 x magnification tomograms from two independent cultures for control (n=39 tomograms) and NbLumSyt1 (n=36 tomograms) **(h-w)** 57 000 x magnification tomograms from one culture; NbLumSyt1 (n=11 tomograms).

Quantification of the spatial distribution of presynaptic vesicles revealed that CCV^G^s were positioned closer to the AZ than unlabeled CCVs (**Fig. 2o-p**), indicating that recycling vesicles labelled during spontaneous network activity are repositioned in AZ proximity for preferential reuse. Tomographic sub-volumes containing endocytic intermediates showed gold-labeling within newly forming vesicles at the plasma membrane, indicating active endocytosis (**Fig. 2r-s**). Further comparison of the CCSV^G^s’ proportional representation relative to SVs across tomograms demonstrated the preferential labeling of recycling vesicles by NbLumSyt1-ALFA-tag:NbALFA-Gold complex (**Fig. 2w**). These ultrastructural analyses collectively indicated that the NbLumSyt1-ALFA-tag:NbALFA-Gold complex labels actively recycling SVs with high specificity, providing a robust framework for investigating mechanisms of vesicle trafficking at the synapse.

### Exo- endocytic cycling of SVs monitored with novel NbLumSyt1-reporters

To test whether labeling with NbLumSyt1 affects synaptic function, we analyzed SV exo- and endocytosis in living neurons. To this end, we created fusion proteins in which NbLumSyt1 was fused to the GFP variants e-pHluorin ^24^ (pHluorin) or mOrange both quenched in the acidic interior of SVs fluorescent when exposed at the plasma membrane. To ensure minimal perturbation, miniature excitatory postsynaptic currents (mEPSCs) were recorded after loading neurons with either the NbLumSyt1-pHluorin or a commercial Syt1 luminal antibody and compared to controls. Prolonged loading (60 min) resulted in a small, non-significant increase in mEPSC frequency. In any case, to avoid unnecessary alterations in all subsequent experiments, we decided to use a 30-min loading, since the signal was sufficient, and no changes of activity were observable (**Fig. 3a-c**). Nonetheless, we prompt general prudence, as Syt1 luminal domain antibodies have previously been associated with partial loss-of-function effects ^25^.

**Figure 3:**
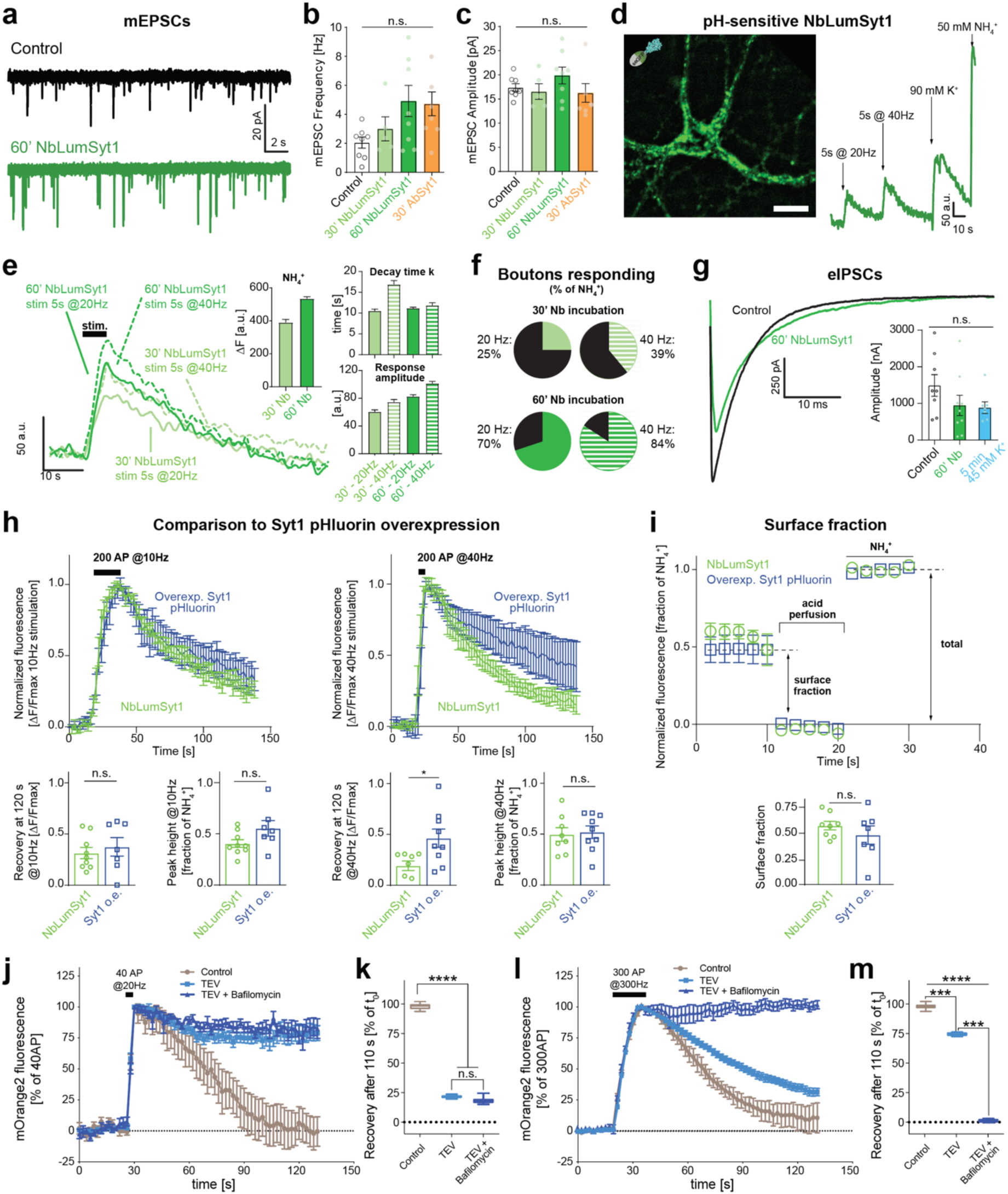
The NbLumSyt1 is a minimally invasive tool to measure synaptic vesicle exo-endocytosis. **a-c**) Miniature excitatory postsynaptic current (mEPSC) measurements in control (no live uptake) and treated neurons with different times of loading either with the NbLumSyt1 directly fused with pHluorin or with a commercial Syt1-luminal antibody (SySy 604.2). When longer incubation times (60m) were used with the nanobody and the antibody, there was a tendency, although not significant, of increased frequency. For this reason, for subsequent experiments (panels h-i), loading was performed for 30 min only. **d**) Live imaging of the NbLumSyt1-pHluorin recycling. Fluorescence amplitude scales with stimulation intensity, indicating more nanobody uptake, followed by florescence decay, suggesting endocytosis of the nanobody. Treatment with ammonium chloride (NH_4_^+^) reveals the total pool of labeled vesicles. **e**) Example traces, with decay time constants (tau) and amplitudes upon electrical stimulation of cells labeled with NbLumSyt1-pHluorin. **f**) Percentage of boutons responding to different electrical stimuli (calculated as % of the total pool of SVs, as defined by NH_4_^+^ treatment). **g**) Evoked inhibitory postsynaptic currents (eIPSC), to study action potential drive GABAergic neurotransmission, control neurons where stimulated 5 min in 45 mM KCl without nanobody. Inset: Quantification of eIPSC amplitude. **h**) Comparison of NbLumSyt1-pHluorin to neurons overexpressing Syt1 pHluorin. The overexpression of Syt1 pHluorin causes slower endocytosis at higher (40 Hz) stimulation frequencies. **i**) Evaluation of the surface fraction by perfusion with an acidic buffer (pH 5). Ammonium chloride perfusion at the end of the experiment is used to reveal the total population of molecules (used for normalization). The surface fraction is unchanged at our Syt1 pHluorin expression levels. **j**-**l**) To study the involvement of the surface population of Syt1, neurons were loaded with a TEV-cleavable version of the nanobody (NbLumSyt1-TEV-mOrange2). After 5 min of incubation with the TEV protease, the protease was washed out. Neurons were then field-stimulated either for 2 s at 20 Hz (**j**) or for 10 s at 30 Hz (**f**) in the presence or absence of the re-acidification blocker Bafilomycin. The graphs summarize the results of 3 independent experiments. In the control condition, as expected, fluorescence is quenched and returns to baseline after endocytosis due to acidification of the endocytosed Syt1 pools. The fluorescence in TEV-cleaved neurons does not recover after mild stimulation (**j,** recovery at 110s quantified in **k**) confirming that under these conditions the pre-existing TEV-cleaved membrane pool is preferentially internalized. In contrast, after more pronounced stimulation, fluorescence quenching is observed, which is explained by the use of Syt1 originating from TEV-protected vesicles that are recruited during recycling (**l**, recovery at 110s quantified in **m**). This can be explained by the depletion of the readily retrievable vesicle pool caused by more pronounced exo-endocytosis and the engagement of the freshly exocytosed Syt1-containing pool (which was TEV-protected and still contained the pH-sensitive mOrange2).

Next, live imaging of NbLumSyt1-pHluorin recycling revealed dynamic changes in fluorescence upon electrical stimulation. A major increase in fluorescence was observed by exocytotic exposure of the previously internalized nanobody to the extracellular surface, followed by a rapid decay due to endocytosis and re-acidification after the stimulus train. Ammonium chloride treatment, which neutralizes the pH across all compartments including SVs, allowed us to estimate the total pool of labeled vesicles (**Fig. 3d**).

Analysis of fluorescence decay kinetics demonstrated consistent tau values across conditions, supporting the tool’s utility in quantifying vesicle dynamics (**Fig. 3e**). The high proportion of boutons responding to increasing electrical stimuli further confirmed the reliability of this labeling method (**Fig. 3f**).

Comparisons of NbLumSyt1-pHluorin-labeled neurons to those overexpressing Syt1-pHluorin highlighted key differences: Syt1-pHluorin overexpression slowed endocytosis during high-frequency (40 Hz) stimulation (**Fig. 3h**). Perfusion with acidic buffer, which selectively quenches surface-exposed pHluorin, revealed that the size of the plasma membrane pool after exocytosis was unchanged when compared to neurons overexpressing Syt1-pHluorin. Together, the data show that labelling of neurons with NbLumSyt1 does not perturb SV recycling kinetics (**Fig. 3i**).

To differentiate between distinct surface-exposed and internalized pools of NbLumSyt1-tagged Syt1, we created modified fusion proteins in which NbLumSyt1 was connected to reporter domains via a linker containing a cleavage site for tobacco etch virus (TEV) protease (*i.e.*, NbLumSyt1-TEV-pHluorin, NbLumSyt1-TEV-mOrange). Efficient cleavage was observed when analyzing the products by SDS-PAGE after TEV-treatment (**Supplementary Fig. 4a**). When neurons were pre-incubated with the mOrange2 variant and then treated with the TEV-protease to remove the fluorescent surface-exposed pool, cleavage was completed within few minutes (**Supplementary Fig. 4b**), similar to that of the purified proteins (**Supplementary Fig. 4c**).

As previously reported ^26^, this approach allows for differentiating between freshly exocytosed and longer-lingering surface pools of Syt1. To this end, neurons preloaded with NbLumSyt1-TEV-mOrange2 were treated with TEV-protease to remove all pre-existing label exposed on the surface under resting conditions. After washout of the protease the neurons were moderately stimulated to selectively exocytose the readily releasable SVs (**Fig. 3j,k**). As expected, fluorescence intensity increased rapidly since the newly exposed intravesicular sensors were protected during protease pretreatment. After the end of the stimulus train, a rapid decay was observed in the untreated samples. However, the decay was much lower in the TEV-pretreated preparation, confirming that upon mild stimulation, the pre-existing plasma membrane pool (rendered non-fluorescent by TEV-pretreatment) is preferentially internalized by compensatory endocytosis ^26^. During longer stimulation trains, however, a much higher proportion of SVs underwent exocytosis, followed by clear endocytosis that in this stimulation paradigm revealed the exo-exocytosis of vesicles and the fact that the readily retrievable pool is limited (**Fig. 3l,m**). Note that in the presence of bafilomycin, an inhibitor of the vesicular V-ATPase, used here as a control, no decay of the fluorescence signal was observed during endocytosis since re-acidification was inhibited ^27^.

### Resting pH estimations with GFP-based nanobody reporters

We utilized the NbLumSyt1-pHluorin and mOrange2 constructs to revisit some of the previous work on the resting pH of SVs in synaptic boutons ^28^. mOrange2 has a lower pK-value than pHluorin, thus allowing to extend the measuring range towards lower pH-values such as those found in SVs ^28^ (**Supplementary Fig. 4**). To this end, neurons were allowed to internalize the reporters, followed by TEV-cleavage to remove all extracellular label. After recording fluorescence, boutons were pH-clamped at defined pH values for calibration, fixed, and immunostained for the vesicular glutamate (vGluT and GABA transporters (vGAT) to differentiate between excitatory and inhibitory boutons (**Fig. 4**). Significant differences were observed, with glutamatergic neurons showing a lower resting pH than the GABAergic neurons (**Fig. 4d,e**), in agreement with previous results ^28^.

**Figure 4:**
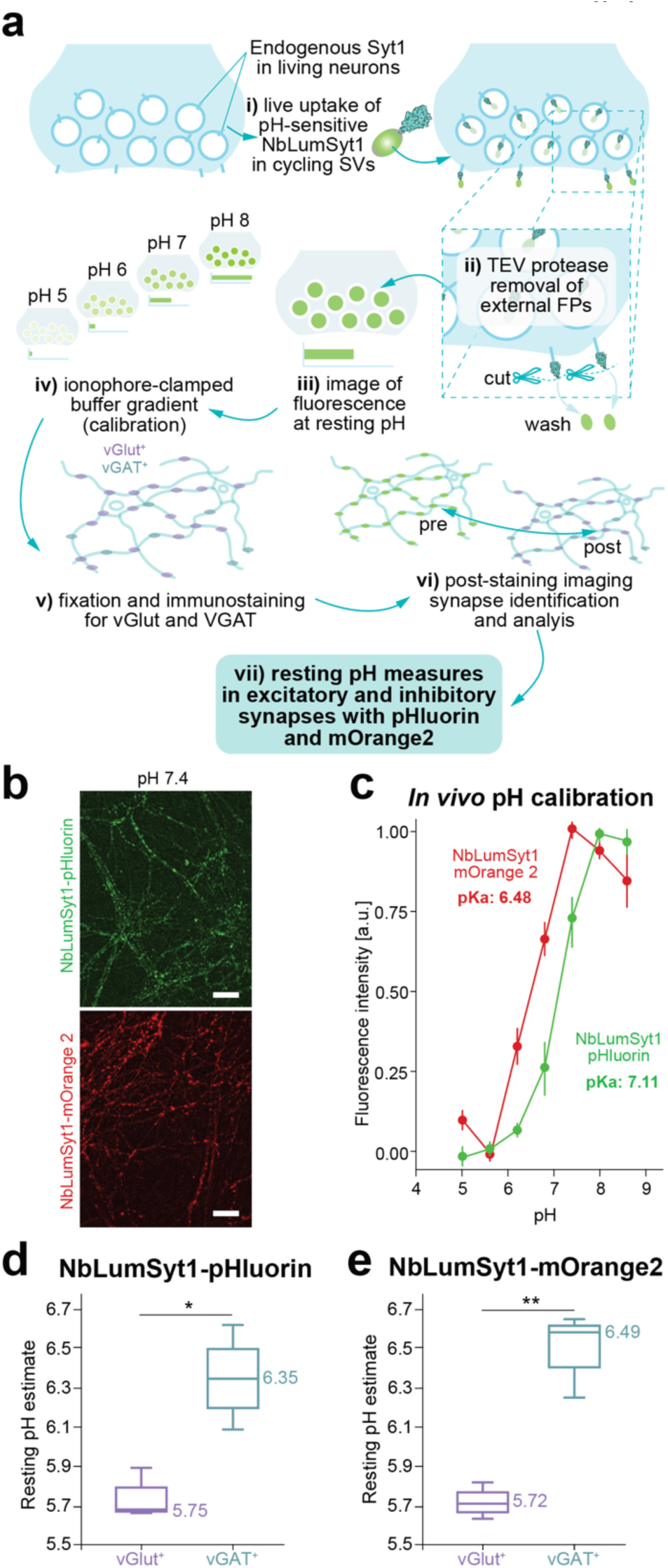
Estimation of resting pH using NbLumSyt1-pHlourin and NbLumSyt1-mOrange2 probes in rat hippocampal neurons. **a**) Scheme of the experimental design for estimation of the resting pH n SVs. The resting pH of either excitatory or inhibitory boutons was estimated similarly to what previously described ^28^ albeit with minor modifications (see methods). Neurons were labeled with either the NbLumSyt1-pHluorin or the NbLumSyt1-mOrange2 (**b**) in both cases containing a TEV-recognition site for the TEV protease. The pH-sensitive sensors were cut from the surface pool by treating cells with the TEV protease for 3 min, which ensured efficient removal of the pH-sensitive fluorescent proteins (see also **Supplementary Fig. 4 a-c**). The image was then captured and reflected the “resting state”. Neurons were treated with buffers containing ionophores that were adjusted to increasing pH values, and images were acquired at different pH steps. At the end of the experiment, the sample was fixed and immunostained with antibodies specific for vGlut and vGAT, the same region was located again (post-hoc) and re-imaged. All images were aligned and analyzed. The pKa values estimated from in vitro (**Supplementary Fig. 4 d-f**) and in vivo (**c**) allowed the generation of calibration curves that were used for calculating the resting pH values (see methods). The pH-response of the probes was titrated. In accordance with previous reports ^28^, the pKa of mOrange is lower than that of pHluorin (**Supplementary Fig. 4 d-e**), extending the measuring range towards lower pH-values. **d,e**) The values obtained by the two probes were similar and revealed a significant difference between excitatory and inhibitory boutons (NbLumSyt1-pHluorin (**d**), NbLumSyt1-mOrange2 (**e**)). Unpaired Student’s t-test, N=3. * = p ≤ 0.05, ** = p ≤ 0.01. Note that numbers in the graphs indicate average values.

### Live-cell proteomic mapping of Syt1 surface interactions using NbLumSyt1-APEX2

To perform live-cell mapping of Syt1 surface interactions with other proteins and potentially identify novel Syt1 interaction partners/modulators, we developed a version of the nanobody conjugated to APEX2 ^29^. This enzyme enables efficient biotinylation of proteins in close proximity of a protein of interest, in our case Syt1, during live-cell uptake.

Representative experiments with the NbLumSyt1-APEX2 show that biotinylation only occurs when all reaction components, including H_2_O_2_, are present, as revealed by fluorescent streptavidin staining (**Fig. 5a**). WB analysis confirmed robust biotinylation under these conditions (**Fig. 5b**). Moreover, *in situ* proximity labeling performed with NbLumSyt1-APEX2, coupled with deposition of electron-dense material visible in EM, revealed labeled SVs, demonstrating the specificity of this approach (**Fig. 5c**). Having established efficient protein biotinylation and targeting, we performed protein identification following labelling and streptavidin bead pull-down. Proteome analysis by liquid chromatography followed by Mass Spectrometry (LC-MS/MS) showed enrichment of Syt1 (positive control) and other interactors (**Supplementary Table 1)**. Controls using neurons without nanobody uptake or labelled with an NbALFA-APEX2 demonstrated minimal nonspecific biotinylation thereby validating the specificity of the approach (**Fig. 5d**). Gene ontology (GO) analysis of the candidates enriched in the NbLumSyt1-APEX2 samples clearly highlighted synaptic and membrane-associated components, consistent with the expected localization of Syt1 interactors (**Fig. 5e**). Interestingly, among the identified interactors, the ciliary neurotrophic factor receptor (Cntfr) was the most enriched hit, alongside additional proteins that may be the subject of future studies (**Fig. 5f**). Cntfr is a cell surface receptor involved in neuroprotection and synaptic maintenance ^31^, whose potential partnership with Syt1 could regulate SV dynamics, specifically changing vesicle release or recycling efficiency at synaptic terminals.

**Figure 5:**
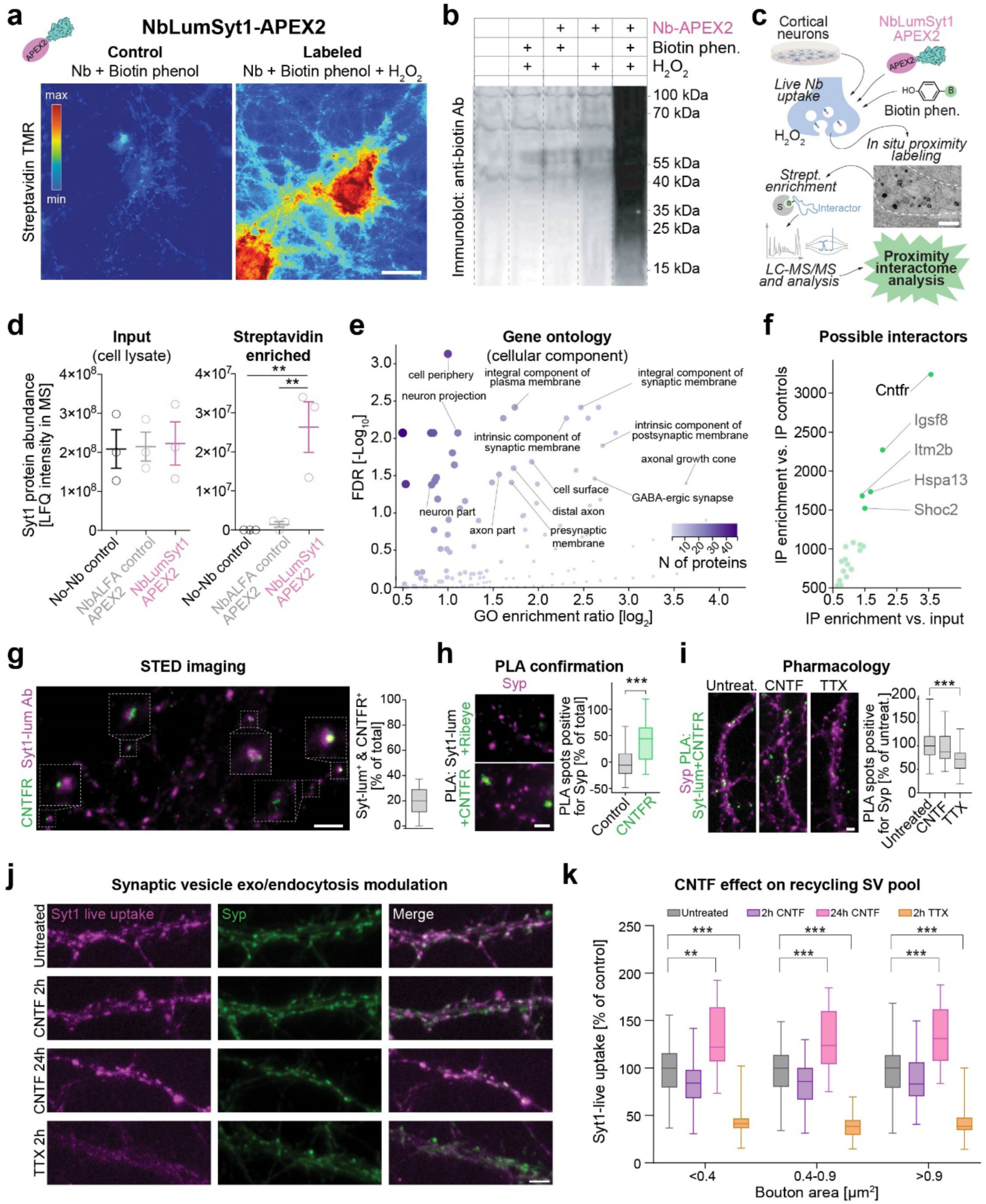
Syt1 is found in close proximity of the ciliary neurotrophic factor receptor (Cntfr) and regulates synaptic vesicle dynamics. **a-c**) The NbLumSyt1-APEX2 allows efficient biotinylation of proteins in the proximity of Syt1 upon live uptake in hippocampal neurons to facilitate live-cell proteomic mapping. Representative images of neurons upon uptake of the NbLumSyt1-APEX2 where biotinylated proteins are revealed with fluorescent Streptavidin in **a**. In the absence of H_2_O_2_, only the few endogenous biotinylated proteins are observable. In the presence of all components, the reaction occurs efficiently, as also revealed by western blot of labeled neurons (**b**). To identify the interactors of Syt1, in situ proximity labelling was performed with the NbLumSyt1-APEX2 (**c**). The electron microscopy image in the scheme is an example of the labelled vesicles, as reveled upon photoconverting 3,3-diaminobenzidine (DAB) into a stable, electron microscopically visible dark product. **d**) Protein intensities measured with LC-MS/MS in input and upon enrichment of the biotinylated proteins. For controls neurons without nanobody or neurons where the anti-ALFA-Nb was provided in the medium of neurons are used. Note that, since the primary neurons do not express the ALFA-tag, this control will allow to reveal the effect of the unspecific biotinylation of the membranes occurring during the labelling period. Note that Syt1, as expected, is efficiently biotinylated and enriched upon IP with streptavidin beads. For details concerning this type of experiments and the respective analysis, refer to the methods. **e**) Summary of the gene ontologies (GOs; cellular components) for the proteins biotinylated upon live uptake of the NbLumSyt1-APEX2 (for a detailed list see **Supplementary Table 1**). As expected, synaptic components and membrane GOs are over-represented. **f**) Possible interactors identified upon live-cell proteomic mapping and enrichment vs. input and vs IP control. Cntfr was found to be the most enriched candidate, together with other proteins that could be studied in future works. As an additional interesting candidate, the integral membrane protein 2B (Itm2b) stands out since it plays a role in vesicle trafficking, is linked to neurodegenerative diseases involving synaptic dysfunction, and may regulate SV recycling and neurotransmitter release^30^. **g**) Super-resolution stimulation emission depletion (STED) imaging reveals that ∼20% of boutons labeled with live uptake are also positive for Cntfr. Note that in this case, for cross-validation purposes, the live uptake was performed with the Syt1-luminal antibody. **h**) Proximity ligation assay (in situ PLA), using antibodies against the luminal portion of Syt1 and anti-Cntfr, confirms close proximity for these two proteins. As a control, a primary antibody against a protein not expressed in hippocampal neurons (Ribeye) was used. **i**) Blocking the network activity of primary hippocampal neurons with tetrodotoxin (TTX) decreases the in situ PLA signal between Syt1 and Cntfr. Stimulation with the ligand of Cntfr (Cntf; 8 nM) does not change the PLA signal between Syt1 and Cntfr. **j-k**) Stimulation of neurons with Cntf, increases SV exo-endocytosis following 24h incubation.

To further characterize the possible local interaction of Syt1 and Cntfr, we performed super-resolution stimulation emission depletion (STED) imaging on neurons co-staining for the two proteins and using an antibody against the Syt1-luminal domain (Syt1-lum Ab) for cross-validation purposes. These experiments showed that ∼20% of synaptic boutons positive for Cntfr were also positive for live uptake with Syt1, confirming that the two molecules indeed are found in close proximity (**Fig. 5g**). To confirm their direct local interaction, a proximity ligation assay (PLA) with Syt1-lum and Cntfr was established and revealed close proximity, with a minimal signal detected when a non-neuronal protein was used as a negative control (**Fig. 5h**). To further test possible functional crosstalk between the two molecules, we performed experiments modulating the activity regimens of our neurons. Network activity blockade with tetrodotoxin (TTX) decreased the PLA signal between Syt1 and Cntfr, showing that inhibition of neuronal activity leads to a decrease in the interactions of the two proteins. Importantly, direct stimulation with the Cntfr natural ligand, Cntf, for 2h did not alter the PLA signal, indicating that Cntf binding does not significantly affect Syt1-Cntfr proximity at short time points (**Fig. 5i**). However, prolonged stimulation with Cntf (24 hours) led to an increase in SV exo-endocytosis, revealing a role for Cntfr in regulating synaptic activity (**Fig. 5j-k**).

### Nanoscale organization of endogenous surface-stranded Syt1 using NbLumSyt1-Halo reveals two populations of molecules with distinct dynamics

Nanobodies offer several advantages for diffraction-unlimited imaging ^32^. This has recently allowed the characterization of the surface clustering of Syt1 following exocytosis and established a novel role in coordinating the binding and SV targeting of botulinum neurotoxin type-A ^33^. To track endogenous Syt1 behavior in live conditions following exocytosis and avoid perturbations due to overexpression, we performed single-molecule imaging of surface endogenous Syt1 in live hippocampal neurons with universal Point Accumulation for Imaging in Nanoscale Topography (uPAINT) ^34,35^. Labeling of Syt1 was achieved via incubation with pre-conjugated complexes: either NbLumSyt1-HALO bound to the Halo-ligand JF549 or NbLumSyt1-pHluorin pre-complexed with the anti-GFP nanobody conjugated to Atto647N. Imaging was done using highly inclined and laminated optical microscopy (HILO) to increase signal-to-noise discrimination. For comparison, neurons overexpressing Syt1-pHluorin were imaged in identical conditions using the anti-GFP nanobody Atto647N (**Fig. 6a**) ^36^. This experimental setup allowed the detection and tracking of single endogenously and exogenously expressed Syt1 molecules with high temporal resolution (20 ms), providing super-resolved images of single-molecule trajectories and diffusion coefficients (**Fig. 6b**). The map of diffusion coefficients showed heterogeneous mobility, with colder colors representing lower mobility, while average intensity maps, which indicated localization densities, showed distinct regions of lower (cold colors) and higher detection densities (white color; **Fig. 6b**).

**Figure 6:**
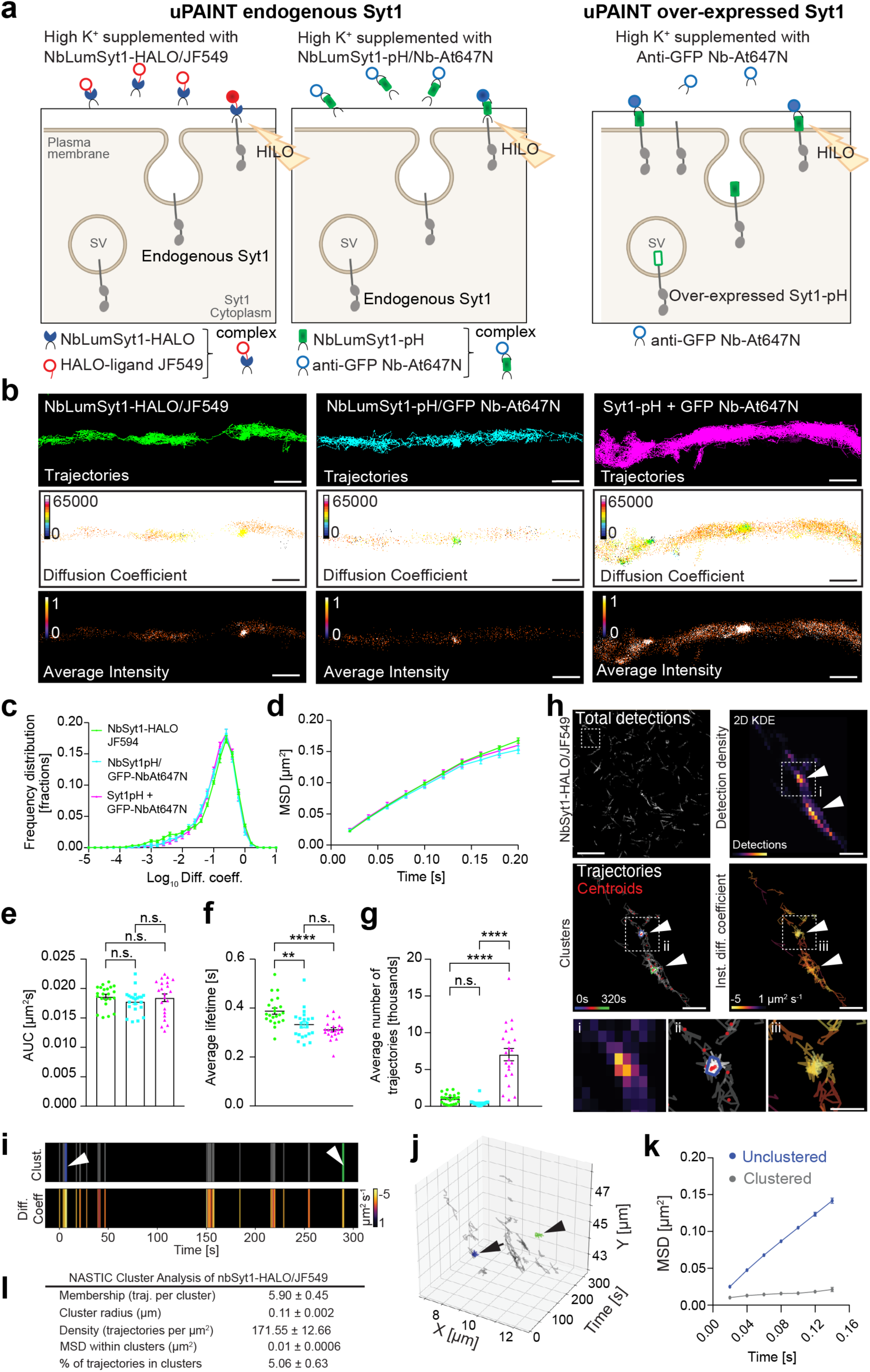
Live cell single molecule imaging of endogenous Syt1 reveals two populations of molecules that differ in their displacement dynamics. **a**) Scheme of single fluorescent molecule detection in live hippocampal neurons using uPAINT. To detect and track single molecules cells were incubated with complexes conjugated prior to imaging the NbLumSyt1-Halo-Tag with the Halo-ligand JF549 or the NbLumSyt1-pHluorin with the anti-GFP Nb-At647N (NbLumSyt1-pH/Nb-At647N). Neurons at 21-22 days in vitro (DIV) were washed and stimulated in high K+ buffer supplemented with 1 nM complexes, engineered to bind endogenous Syt1, and imaged in HILO for 320 seconds at 50 Hz. As a comparison, hippocampal neurons were transfected with Syt1-pHluorin (Syt1-pH) and uPAINT imaging was performed using high K+ buffer supplemented with 1 nM anti-GFP nb-At647N, which specifically binds to the pHluorin moiety. **b**) Super-resolved images of single molecule trajectories, diffusion coefficients, and average intensities in hippocampal neurons over 16,000 frames acquisition. The color scale in the Diffusion Coefficient map ranges from 0 to 1, corresponding to Log_10_ diffusion coefficient detections, and the colder colors in the scale indicate lower mobility. The average intensity map represents localization densities as arbitrary units, with warmer colors indicating higher detection densities. Scale bars, 1 μm. **c-g**) Quantifications of the parameters in the different conditions. The lifetime decreases with larger sizes of the complexes and the over-expression of Syt1 results in significantly higher number of trajectories compared to the endogenous Syt1 trajectories. Statistical analysis of normally distributed data was performed using one-way ANOVA Multiple comparison in g, and for non-normally distributed data, one-way ANOVA Kruskal-Wallis test Multiple comparison was used. Asterisks indicate the following p-values *p < 0.05, ** p < 0.01, and **** p <0.0001. **h**) Representative image showing detections from a 320 s acquisition at 50 Hz of the NbLumSyt1-Halo/JF549 complex imaged by uPAINT in cultured hippocampal neurons. The arrow and arrowhead indicate two separate NbLumSyt1-Halo/JF549 clusters and the boxed areas (i-iii) are magnified. The resulting NASTIC analysis images of 2D kernel density estimation (KDE) of detections with single molecule trajectories, centroids, clusters, and instantaneous diffusion coefficients (Log_10_) from the boxed region are shown magnified in the boxed regions. Trajectories are shown in grey, with each trajectory segment representing the displacement of the detected molecule in 0.02 s. The centroid of each trajectory is indicated with a red dot. Colored convex hulls indicate the extent of the detections associated with clustered trajectories as determined by NASTIC. **i**) The top panel show the 2D temporal clustering NbLumSyt1-Halo/JF549 (arrow and arrowhead point to the timeframe of blue and green clusters shown in **h**) over 320 seconds and the lower panel the respective diffusion coefficient mobility of the trajectories, indicating a lower mobility of NbLumSyt1-Halo/JF549 within clusters that when unclustered. Scale bars, 1 μm. **j**) 3D presentation of the spatiotemporal clustering of NbLumSyt1-Halo/JF549 where the blue and green clusters from c and f are indicated. **k**) The graph shows the quantification of mean square displacement (µm^2^ s^−1^) of NbLumSyt1-Halo/JF549 within clusters and outside of the clusters (unclustered). MSD curves measure the average mobility with respect to time of all the trajectories observed. **l**) Biophysical properties of NbLumSyt1-Halo/JF549 clusters (n = 10 neurons). Numbers represent average values ± SEM.

Quantification of single-molecule mobilities using the three labelling techniques revealed similar frequency distributions of diffusion coefficients (Log_10_D), mean square displacement (MSD) and area under the MSD curve (AUC) values (**Fig. 6c-e**), while the fluorescent lifetime of labeled complexes decreased with larger-sized labeling tools, which may reflect certain limitations of bulkier affinity tools (**Fig. 6f**). As expected, neurons overexpressing Syt1-pHluorin showed a significantly higher number of trajectories when compared with those labeled by endogenous Syt1. Single-molecule detections and tracks of NbLumSyt1-HALO/JF549 on the membrane of hippocampal neurons showed distinct confinement areas (**Fig. 6h**; Total detections). The spatiotemporal clustering ^37^ of endogenous Syt1 molecules was further interrogated using two-dimensional kernel density estimation (2D KDE) to identify trajectories associated with Syt1+ clusters (**Fig. 6h**). The Syt1^+^ trajectories detected in clusters displayed lower mobility compared to the unclustered molecules, as evidenced by their reduced diffusion coefficients and confinements (**Fig. 6i**). Visualization of the spatiotemporal clustering ^37^ of the tracked Syt1^+^ surface molecules revealed multiple immobilizations zones detected concomitantly with more mobile trajectories over time (**Fig. 6j**) as recently described ^38,39^. Quantifying the MSD of Syt1 molecular dynamics in these spatiotemporal clusters exhibited significantly reduced mobility (**Fig. 6k**). Some areas showed clusters characterized by lower estimated diffusion coefficients associated with reduced mobility. In contrast, the unclustered molecules had higher mobility (**Fig. 6k**), revealing two populations of surface molecules, one probably engaged in more complex interactions (clustered) and one less restrained (unclustered). The quantification of biophysical properties of NbLumSyt1-Halo/JF549 nanoclusters are shown in **Fig. 6l**. This nanoscale analysis clearly identified two subpopulations in Syt1 molecules with distinct cluster association based on their displacement dynamics providing insights into the functional nano-organization of SV components following exocytosis on the plasma membrane. The functional significance of these surface clusters has recently been revealed as Syt1, polysiologanglioside and SV2A forming tripartite surface nanoclusters that act as a receptor for botulinum toxin type-A ^33^. Combined with our demonstration that Syt1 overexpression alters the rate of endocytosis at high stimulation rate, our data suggest that altering Syt1 synaptic abundance impacts the maintenance of synaptic function.

To further investigate the use of NbLumSyt1 in another live-imaging diffraction-unlimited modality, we used MoNaLISA (Molecular Nanoscale Live Imaging with Sectioning Ability) imaging ^40^, parallelized RESOLFT approach tailored to the dynamic organization of molecules in living cells. In this imaging modality SVs were live-labeled with the NbLumSyt1-rsEGFP(N205S) in neurons. Through the combination of extended light patterns, MoNaLISA enhanced the spatial resolution to nanoscale clusters as small as 65 nm with inter-cluster distance of approximately 130 nm over an extended field of view in at a frame time of 2 seconds (**Supplementary Fig. 5a**). Comparisons between the wide-field and MoNaLISA images illustrated the resolution gained (**Supplementary Fig. 5b, c**). Time-lapse imaging of NbLumSyt1-rsEGFP(N205S)-labeled vesicles demonstrated marked structural rearrangements, even at 1 min separation (**Supplementary Fig. 5d, e**). Quantification of single clusters showed dynamic changes in their size, ellipticity, and intensity (**Supplementary Fig. 5f**). These findings support that SV organization rapidly changes at the nanoscale. Further, dual-color imaging of NbLumSyt1-rsEGFP(N205S) together with the inhibitory synapse marker nanobody anti-vGAT-Cy3 showed enrichment of the labeled clusters at inhibitory synapses (**Supplementary Fig. 5g-i**). Quantitative comparisons showed that inhibitory synaptic clusters were larger in area and more elliptical in shape and had higher mean fluorescence intensity compared to their excitatory counterparts. Overall, these features indicate denser and more organized vesicle distribution at inhibitory synapses, consistent with functional specialization in these synaptic domains ^41^, and highlight the usefulness of live-imaging diffraction-unlimited modalities.

### Translational application of NbLumSyt1 for real-time monitoring of synaptic vesicle recycling in human neuronal systems

To showcase the translational potential of NbLumSyt1, we characterized the performance of our nano-binding tools in human neurons. NbLumSyt1 was specific for the binding of human Syt1 in human iPSC-derived hypothalamic-like neurons and had the potential to monitor SV recycling under physiological conditions. Live staining with NbLumSyt1 followed by fixation and subsequent immunoassaying for endogenous Syt1 and the neuronal marker Beta-III-tubulin (TuJ) confirmed these cells’ neuronal identity and synaptic pattern. Pre-treatment with TTX strongly reduces activity-dependent staining, validating that most labeling had been driven by neuronal activity. A basal level of staining persisted reflecting surface binding and spontaneous vesicle recycling occurring independently of evoked activity (**Fig. 7a**). Quantification revealed that stimulation during labeling strongly increased the overall luminal intensity of Syt1 staining. A positive correlation between the luminal and total Syt1 staining levels further confirmed the dependence of the probe on activity upon stimulation conditions. In agreement with the sensitivity to physiological synaptic activity, NbLumSyt1 staining levels were significantly reduced when cells were treated with TTX (**Fig. 7b**).

**Figure 7:**
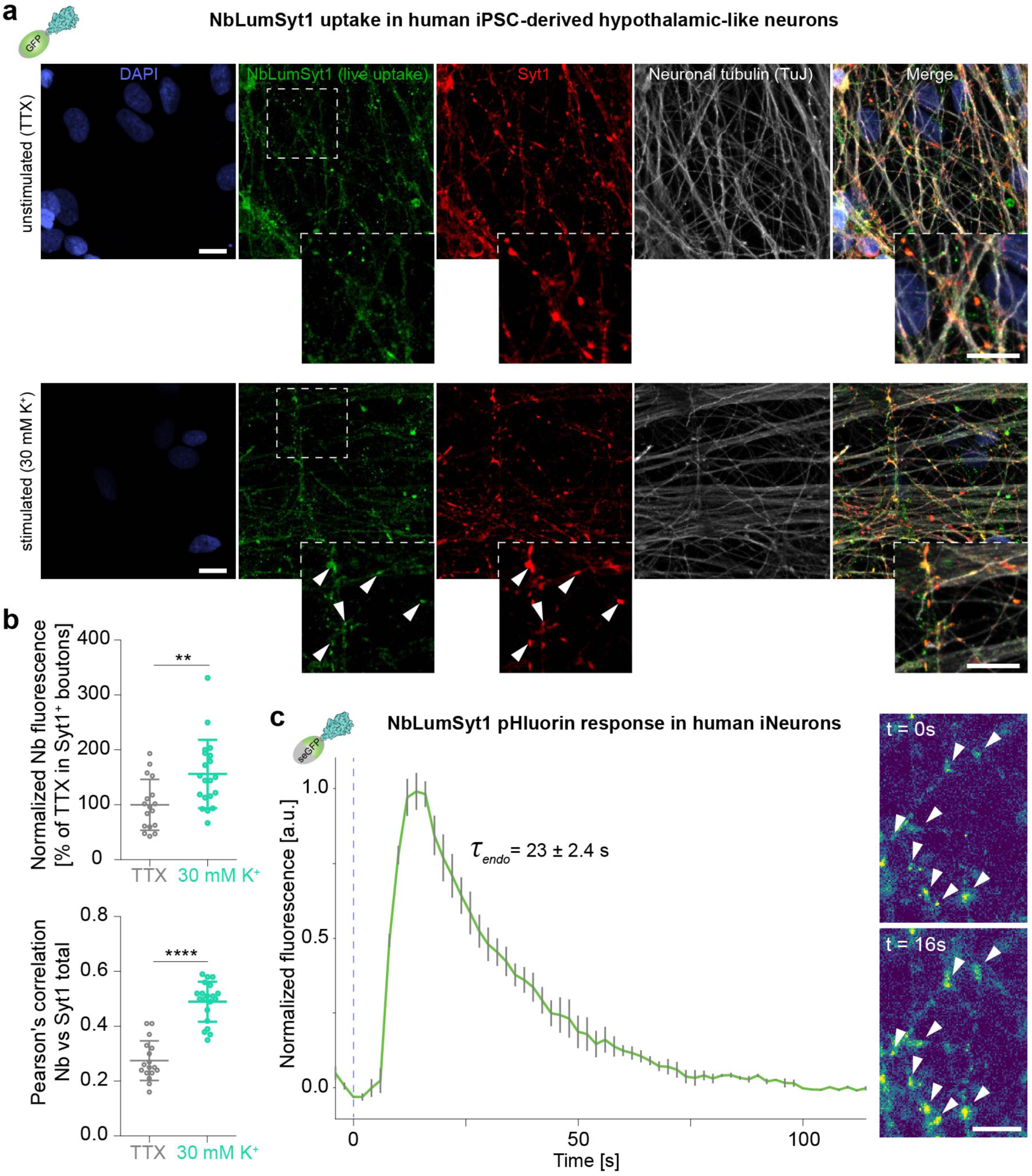
The NbLumSyt1 specifically binds human Syt1 in living cells and can be used for recycling assays in human iNeurons. **a**) Human iPSC hypothalamic-like neurons were stained live with the NbLumSyt1 and after recycling were fixed and immunoassayed against the endogenous Syt1 and the neurospecific BetaIII tubulin marker (Tuj) to ascertain the neuronal commitment of these neurons. In cells pretreated with TTX during labelling. Scale bars, 10 μm. **b**) Quantification indicates that increased stimulation during labelling increases both the labelling level (upper panel) and the correlation between the luminal Syt1 and the total Syt1 (lower panel). A certain amount of staining in TTX-treated cells is due to a combination of surface staining and a certain degree of activity-independent spontaneous recycling. Unpaired Student’s t-test: ** = p < 0.01; **** = p < 0.0001. **c**) Synaptic vesicle recycling measured in human iPSC-derived neurons (iNeurons) monitored with the pH-sensitive version of the NbLumSyt1. (Left) Averaged normalized pHluorin fluorescence traces from iNeurons (6-8 weeks in culture, N=3) stimulated with 200 action potentials (40 Hz, 5s) at physiological temperature (37t °C). the blue segmented line indicates the begin of the stimulation. (Right) Representative images showing the sensor intensity before (t = 0s) and directly after (t = 16s) the electrical stimulation. Scale bar, 5 μm.

To label SVs directly in human iNeurons ^42^ and thereby measure SV recycling, the NbLumSyt1-pHluorin was used. Stimulation with 200 action potentials at 40 Hz elicited a dynamic fluorescence response that captured the temporal progression of exocytosis followed by endocytosis at physiological temperature. Averaged fluorescence traces demonstrated the reliability of NbLumSyt1-pHluorin in quantifying SV dynamics in real-time. Representative images before and after stimulation showed the resolution and sensitivity of the tool in tracking these processes at the synaptic level (**Fig. 7c**). These findings highlight the usefulness of NbLumSyt1 as a reliable molecular probe to study SV recycling mechanisms in human neuronal systems.

## Discussion

The development of NbLumSyt1 represents a significant advance within the synaptic pathophysiology toolkit allowing flexibility and future applications for human studies to interrogate SV recycling and synaptic physiology in a clinically relevant context. This nanobody-based approach targets the luminal domain of Syt1, a central regulator of SV exocytosis and endocytosis, in a highly specific, minimally invasive and versatile manner. Although Syt1 has long been pointed out in previous studies to play a crucial role in calcium-dependent neurotransmission and SV recycling ^7^, restrictions in the available arsenal of tools have hampered real-time and high-resolution visualization of these processes, especially in human neurons. NbLumSyt1 overcomes these challenges by offering a means of combining fluorescence-based live imaging, ultrastructural studies, and proteomic analyses in a future clinically oriented study of synaptic physiology.

The live-cell imaging applications of NbLumSyt1, especially coupled with either HALO or pHluorin tags for the acquisition of spatiotemporal dynamics in the process of SV recycling, highlight the flexibility of this tool. Such experiments revealed spatial distributions of SV pools and their dynamics during recycling. Our results align well with the previously reported heterogeneity and dynamic nature of SV pools ^11,38,39^. Notably, the high signal-to-noise ratio achieved with NbLumSyt1-HALO and the multiplexing capabilities provided by an extensive palette of available HALO ligands ^43^ underlines its potential to detect subtle changes in SV behavior under physiological and pathological conditions. Furthermore, the fluorescence-based pHluorin assays offer simple, convenient and minimally invasive means to estimate intravesicular pH, an important parameter controlling neurotransmitter loading and release ^24^. These tools are an essential step forward to bridge the long-standing gap in measuring SV recycling dynamics without disrupting synaptic function ^44^. While previous studies have raised debate about the role of the luminal region of Syt1 in glycosylation and vesicle targeting ^45,46^, our work seems to suggest that the nanobody has no direct influence on these processes. Although no functional effects were observed in our study under our conditions, previous observations with antibodies ^25^ suggest that potential effects should be evaluated on a case-by-case basis.

Ultrastructural studies of NbLumSyt1-ALFA-tag further validated their specificity and functional relevance. The EM labeling on active SV recycling pathways has shown preferential vesicle recruitment with distinct molecular signatures in support of selective vesicle reuse during synaptic activity. This study thus offers an additional tool to support nanoscale structural analysis and functional investigations by combining advanced EM with nanobody-based labeling modalities for nanogold staining.

Another critical approach is the development of proteomic mapping by employing NbLumSyt1-APEX2 to enable probing of novel interacting proteins involved in regulating SV dynamics. Of these, the interaction of Syt1 with Cntfr points out a novel pathway that could fine tune synaptic plasticity and function. Future experiments will need to clarify the mechanistic regulation of this interaction, and how it integrates with known signaling cascades to modulate synaptic vesicle recycling and neurotransmitter release efficiency

Single-molecule and nanoscale imaging using NbLumSyt1 tools further underline their versatility. Distinct Syt1 populations with different mobility and clustering properties identified in this work align well with previous findings ^11,38,39^ and offer new insights into the organization and dynamics of SV proteins. Moreover, our study suggests that overexpression of Syt1-pHluorin can affect single-molecule mobility and endocytosis. These overexpression-related artifacts are circumvented by using the NbLumSyt1 that selectively binds the endogenous protein. These single-molecule approaches complement advanced imaging techniques, including MoNaLISA /RESOLFT microsxopy ^40,47,48^, which allow live imaging nanoscale description together with other technologies recently developed such as MINFLUX nanoscopy and SUM-PAINT for extensive multiplexing of synaptic targets ^41,49^. By offering a nanobody-based methodology compatible with single-molecule imaging, this study extends the methodology repertoire when studying synaptic nanostructure and function at unprecedented resolution.

The NbLumSyt1 is of particular translational interest. Such a demonstration of effective labeling within human iPSC-derived neurons ensures its applicability within human systems opening the door for the investigation of the SV recycling of more relevant synaptic neurodevelopmental and neurodegenerative disease models. Given the central role that SV dynamics play in both synaptic function and dysfunction ^50^, this could be a very valuable tool in the study of diseases such as Alzheimer’s disease, Parkinson’s disease, and autism spectrum disorders, where disturbances in SV recycling have been implicated. Importantly, NbLumSyt1 is further compatible with a wide range of experimental platforms, including live imaging, proteomics, and nanoscale analyses, thereby further increasing its versatility for basic and translational applications. The various approaches described in the NbLumSyt1 toolkit open the way to investigate neurodevelopmental processes including synaptic maturation and pruning. Real-time imaging of SV recycling in developing neurons may reveal mechanisms underlying synaptic refinement during critical periods of brain development ^51^. Such studies assume particular significance considering the emerging links between disrupted synaptic development and conditions such as schizophrenia and intellectual disabilities ^52,53^. Monitoring SV dynamics in neurodegenerative diseases, especially in the early asymptomatic phases ^54^, may provide insights into the early synaptic dysfunction that could point toward biomarkers or therapeutic targets for conditions such as frontotemporal dementia and Huntington’s disease.

Future applications for NbLumSyt1 might include combining this tool with optogenetic or chemogenetic approaches ^55^ to manipulate and monitor the dynamics of SVs in real-time. The adaptation of NbLumSyt1 will thus allow insights into synaptic function within intact neural circuits. Furthermore, the use of NbLumSyt1 in combination with advanced computational modeling will allow for quantitative analyses of SV recycling kinetics and their regulation under various physiological and pathological conditions. Beyond neuroscience, the experimental workflow that we have developed can inform broader uses, such as drug screening against synaptic proteins or studies of mechanisms in vesicle trafficking in other cellular contexts. In conclusion, the NbLumSyt1 has a significant transformative potential towards the study of SV recycling by presenting a versatile, powerful platform for elucidating the molecular and functional basis of synaptic transmission. Its high specificity, non-invasiveness, and good adaptability make it stand out as a critical new tool for further advancements in the investigation of synaptic physiology and disease alterations.

## Online Methods

### Discovery of anti-luminal Syt1 nanobodies

Immunization and initial screenings were performed in a fee-for-service manner by NanoTag Biotechnologies GmbH (Göttingen, Germany). The intraluminal sequence of Syt1 from *Rattus norvegicus* was expressed and purified from *E. coli*. Two alpacas received six injections with 500 μg of total protein administered once a week. Two weeks after the last immunization, a final boost injection of 500 μg of antigen was used; and blood was drawn on days 3 and 5 post-boost. PBMCs were isolated through a Ficoll gradient. Total RNA was extracted using the Qiagen RNA extraction kit. Using reverse transcription by SuperScript IV, total RNA extracted from PBMCs was used to make cDNA. The final PCR product was verified on a 1.5% agarose and then cloned into a phagemid using Gibson assembly. The resultant phagemid was electroporated into bacteria TG1 for the generation of libraries. The transformed bacteria were grown in 2YT medium with appropriate antibiotics, and the consequent library was aliquoted and stored at −80 °C. The library was panned three rounds against the purified, biotinylated Syt1 antigen. M13KO7 helper phages infected the library to display nanobodies on phages. Phage precipitation using PEG-8000 and filtering purified phages after an overnight culture was performed. Dynabeads MyOne Streptavidin C1 captured the biotinylated antigen. Elution of phage-bound antigen was used in subsequent rounds of selection. Finally, single clones were first characterized by ELISA using the antigen immobilized on a 96-well plate. Positive clones were sequenced, and candidates were produced for initial validations.

### Nanobody expression and purification

Selected nanobody clones were expressed in NEB Shuffle Express T7 *E.coli* (C3029) was transformed and grown overnight. A single colony was picked and grown in TB medium in the presence of 50 µg/mL Kanamycin until the optical density at 600 nm (BioPhotometer 6131 Spectrometer, Eppendorf AG, Hamburg, Germany) reached 0.8. Expression was then induced with 0.3 mM isopropyl-β-D-1-thiogalactopyranoside (IPTG) and the cells were incubated for 14-16 h at 25 °C and 120 g in a shaker (Infors HT, Switzerland).

Nanobodies were purified using Nickel binding affinity chromatography using the protocol adapted from ^23^. Bacterial cells were centrifuged for 25 min at 4,000 g and 4 °C (rotor: H-12000, Thermo Fisher Scientific, MA, USA). The pellet (15 g wet weight) was resuspended in a 50 mL resuspension buffer containing protease inhibitors and homogenized with a glass douncer three times manually. Cell resuspension was processed twice with the microfluidizer at 18,000 psi (M110-Microfluidizer, Microfluidics, MA, USA) and centrifuged at 40K g at 4 °C for 30 min (rotor: Ti45, Beckman, California, USA). Meanwhile, Nickel-NTA resin beads (HisPurTM Ni-NTA Resin, #88222, Thermo Fisher Scientific) were equilibrated with a resuspension buffer by centrifuging for a minute at 800 g on a tabletop centrifuge at 4 °C. This was repeated three times to equilibrate the beads well. Cell supernatant was collected after the centrifugation and rotated with equilibrated beads at 4 °C for 2-3 hours. The supernatant bead suspension was then run over 50 mL Gravity flow column (Econo-column® Bio-Rad Laboratories, Hercules, CA) and the flowthrough (FT) was collected. Beads were washed three times with 50 mL Wash Buffer and once with the Cleavage buffer. After excessive washing, beads were transferred in a 10 mL cleavage buffer in a 15 mL falcon tube. 1 µM of Ulp1 (The yeast homolog of SUMO protease, purified in Dr. Alex Stein’s lab, MPI-BPC) was added and incubated for another hour at 4 °C on constant rotation. Beads were poured back into the column and flowthrough was collected. They were washed once again with 10 mL Cleavage Buffer and remaining eluted protein was collected and 5mM DTT was added. Nanobodies were further concentrated up to 5 mL using 3 kDa MWCO VivaSpin concentrator (Sartorius Stedim Biotech, Göttingen, Germany). Nanobodies were additionally purified using ion-exchange chromatography on a MonoQ 10/100 GL column (GE Healthcare Life Sciences, Pittsburgh, PA) using an ÄKTA system (GE Healthcare Life Sciences, Pittsburgh, PA). ÄKTA buffers A and B were filtered (0.2 µm membrane filter, Sartorius Stedim Biotech, Göttingen, Germany) and degassed. The column was equilibrated with ÄKTA buffer A and a protein sample was injected into the column. Elution was performed with a bdSUMO protease on column ^56^, and the purity of the eluted proteins was assessed via SDS-PAGE.

### Animals and cell culture

In most cases, neurons were cultured as described previously with minor modifications^57^. Briefly, postnatal day zero (P0) Wistar pups were sacrificed by decapitation, brains were extracted, and hippocampi were dissected and put in an enzyme solution containing Papain for 30 min. Hippocampi were then transferred to the inactivation solution. Digested tissue pieces were triturated using fire-polished glass pipettes, first with a bigger hole and then with a fresh fire-polished pipette with a smaller hole, until there was a smooth, homogenous solution. The cells were passed through a 0.22 µm cell strainer, after which they were centrifuged at 5000 g for 5 min. The supernatant was sucked off carefully and the cell pellet was suspended in serum media. Cells were counted using a hemocytometer (Neubauer, Paul Marienfeld GmbH, Germany) manually. 20,000 cells/cm3 were added in wells of a 12 well plate containing Poly D Lysine (PDL) coated coverslips already containing neuron culture medium. Cells were grown in a 37 °C and 5 % CO_2_ incubator and used for imaging between 14-18 DIV. For electron microscopy experiments neuronal monolayer cultures from P0 C57/BL6J mice were cultured onto astrocyte feeder layers as previously described ^58,59^. Briefly, the astrocytes were grown on carbon- and PDL-coated 6mm sapphire disks, glued onto glass coverslips using Matrigel, at a density of 50,000/well in 12-well plates. Astrocytes were grown in DMEM-GlutaMax supplemented with 10% FBS and 0.2% penicillin-streptomycin for six days, before adding 80uM FUDR overnight, then being replaced with Neurobasal-A supplemented with 2% B27, 1% GlutaMax and 0.2% penicillin-streptomycin (NB-A full medium, Invitrogen) an hour before plating hippocampal neurons. Hippocampi were prepared in ice-cold HBSS (CaCl_2_-, MgCl_2_-) incubated in papain at 37 °C for 30-60 min on the shaker, triturated and plated out at 100,000/well onto the astrocyte feeder layer. Cultured neurons were vitrified by high pressure freezing at DIV 14.

To compare the properties of NbLumSyt1-pHluorin with overexpressed Syt1-pHluorin, primary dissociated hippocampal cultures were prepared from mouse embryos (C57BL/6J, E17.5). The mouse hippocampi were digested with papain (10 U/ml) in Dulbecco’s phosphate-buffered saline (PBS), rinsed with Minimal Essential Medium containing 10% fetal bovine serum (FBS), and then mechanically triturated to obtain a single-cell suspension. Cells were plated on 25 mm diameter coverslips coated with poly-D-lysine and laminin. Cultures were maintained in Neurobasal medium supplemented with 2% B-27, 0.5 mM L-glutamine, and 1% (v/v) penicillin-streptomycin, with 1 mM cytosine β-D-arabinofuranoside added at 3 days in vitro (DIV) to inhibit glial proliferation. Neurons were transfected at DIV7 using Lipofectamine 2000 (Thermo Fisher Scientific, 11668-019) according to the manufacturer’s instructions. Imaging of these neurons was performed between DIV14 and DIV16.

### BCA protein estimation

Pierce™ BCA Protein Assay Kit (#23225, Thermo Fisher Scientific, MA, USA) was used to estimate the concentration of proteins according to manufacturer’s instructions in a 96 well flat-bottomed plate (Greiner GmbH, Germany). Protein standard buffers (#23208, Thermo Fisher Scientific, MA, USA) were used to plot a BSA standard curve. Absorbance at 650 nm was measured in a plate reader (Tecan Genios Pro, Männedorf, Switzerland) according to the manufacturer’s manual (BCA Protein Assay, 2020) modified from ^60^.

### SDS Schägger Gels

Protein samples were analyzed using SDS-PAGE on Schägger gels ^61^. 10 µg of protein samples were mixed with the loading buffer and run on a 10 % resolving gel and 4 % stacking gel. Gels were run at 60 V until the samples entered the resolving gel and then they were run at 120 V (Bio-rad Laboratories, Hercules, CA). The PageRulerTM prestained protein ladder 10-180 kDa (#26617, Thermo Fisher Scientific, MA, USA) was used as a molecular marker. Gels were stained by bathing them in Coomassie stain and boiling in a microwave for a minute followed by shaking for 10 min at RT. They were then incubated in destaining buffer 1 and again boiled for a minute, followed by shaking for 30 min. Finally, they were destained in the destaining buffer 2 overnight and then scanned on a scanner (HP, California, US).

### Western blotting

Gels were transferred on a nitrocellulose membrane after running on Schägger gels according to ^62^ with the following modifications. A TransBlot Turbo Transfer machine (Bio-Rad Laboratories, Hercules, CA) was used to perform the transfer in 7 min according to the manufacturer’s instructions. The membrane was blocked in 5 % milk in TBST/PBST. Further, the membranes were incubated in the required primary antibody prepared in 5 % milk in TBS/TBST/PBST (depending on the Ab stability, TBS or PBS (with or without Tween) was used) at 4 °C overnight or 1-2 hours at RT. Membranes were then washed with TBST/PBST and then incubated in secondary infrared (IR) dye-labeled antibodies prepared in TBS/PBS for an hour at RT. After three washing steps of 15 min each, fluorescent protein bands were visualized using Odyssey CLx IR Imaging System (Li-Cor Biosciences, GmbH, Germany).

### Immunoprecipitation of Syt1 from WT and KO Syt1 mice brains using Nanobody against the luminal domain of Syt1

Syt1 WT and KO Mice brains were provided by the lab of Prof. Volker Hauke, Berlin, Germany. The protocol for Syt1 immunoprecipitation (IP) was adapted from Abcam (Cambridge, UK) (Abcam, 2010) and modified. Pre-loading GFP tagged Nanobody (Nb) on the beads-20 µL of bead slurry (GFP Selector, # N0130, NanoTag Biotechnologies, Germany)/sample was pre-incubated with 5µL of GFP Nb (100 µM, NanoTag, cat. N0305-Biotin, RRID: AB_3075908) in 75 µL of lysis buffer. ‘No Nb’ control was also prepared in the same way. Bead slurry with Nb was constantly agitated for 30 min at 4 °C on an orbital shaker in the cold room. It was then centrifuged for a minute at 1000 g (ThermoScientificTM HeraeusTM) and the supernatant was carefully removed. 1 mL of lysis buffer was added and rotated for 5 min at 4 °C. Beads were ready to use after centrifuging once more for a minute at 1000 g. Lysate from tissue - Frozen brains were thawed and 300 µL of Lysis Buffer per brain was added in an Eppendorf tube (Sarstedt AG & Co.KG, Germany). Brains were homogenized using a small Teflon homogenizer 3-4 strokes up and down manually. Debris and nuclei were pelleted down by centrifugation at 1000 g (ThermoScientificTM HeraeusTM) for 5 min and the supernatant was used for protein extraction. The supernatant was constantly agitated for 30 min at 4 °C on an orbital shaker followed by centrifugation for 10 min at 12,000 g in a microcentrifuge (ThermoScientificTM HeraeusTM). Tubes were removed and kept on ice and the supernatant was transferred to fresh cold low binding Eppendorf tubes. Protein concentration was estimated using the BCA protein estimation kit as described previously. Pre-clearing the lysate helps reduce non-specific binding and reduces the background. 100 µL of lysate was mixed with naked (not the ones with preloaded Nb) bead slurry for 30 min on an orbital shaker at 4 °C. Beads were sedimented by spinning at 1000 g for a minute and the supernatant was used for the IP. Each sample (50 µL lysate and 20 µL slurry) was incubated at 4 °C for an hour on an orbital shaker. Beads were then sedimented by centrifugation at 1000 g for a minute and resuspended in 1 mL lysis buffer in a fresh Eppendorf tube. It was again centrifuged for a minute at 1000 g and the supernatant was discarded. This washing step was repeated one more time. The beads were finally resuspended in 20 µL of SDS buffer and heated at 95 °C for 5 min. The samples were loaded on an SDS gel and western blotting was performed using Syt1 cytoplasmic (SynapticSystem, cat. #105 011, AB_887832) and GAPDH (ThermoScientificTM, cat. MA5-15738, RRID: AB_10977387) antibodies.

### Fluorometry for mOrange2/pHlourin pH calibration

Fluorometry was performed on Fluorolog®-3 (Horiba Scientific, Kyōto, Japan) using 250 μL glass cuvettes (Starna GmbH, Germany) to use the minimum volume of precious fluorophores and proteins. *In vitro* pH calibration of pHlourin/mOrange2 fused proteins was performed as previously described ^28^ with slight modifications. MES buffer was used for pH values 5, 5.6, and 6.2 and HEPES buffer was used for pH values above 6.8. The concentration of 0.8μM of the two proteins was enough to give a substantial number of counts (>100 K counts) on the Fluorolog. The protein was added in 250 μL of each of the pH clamped buffers in Eppendorf tubes and vortexed briefly. The mixture was put in cuvettes and the emission spectra were recorded. Data were analyzed on Graph Pad Prism.

### Nb binding with HaloTag fluorescent ligands

HaloTag fused with NbLumSyt1 (NbLumSyt1-HALO) was produced and purified at NanoTag Biotechnologies. The NbLumSyt1-HALO and HaloTag fluorescent ligands were mixed in equimolar ratio at a final concentration of 10 µM for 15 min at RT. The complex formation was confirmed using fluorescent imaging. HALO-Tag ligands were purchased from Promega.

### Labeling of cultured neurons with Nb/Ab by spontaneous endocytosis

Labeling of neurons was performed as described before ^63^ although with the following modifications. 200 µL of media from an 18 mm diameter coverslip (Karl Hecht Assistant, Germany) containing cultured hippocampal neurons was taken in an Eppendorf tube and mixed with the required labeled antibody (Ab) or a nanobody (Nb) (5uM final concentration). The mixture was centrifuged at 13.000 g for 10 min at 37 °C to remove the dye aggregates. The coverslip was incubated in the media for 10 min and placed in a 37 °C/5 % CO_2_ incubator to label the recycling pool of SVs. Neurons were then washed fast 3 times with a warm Tyrode’s buffer (124 mM NaCl, 5 mM KCl, 30 mM glucose, 25 mM HEPES, 2 mM CaCl2, 1 mM MgCl2, pH 7.4). Experiments were performed on neurons from 14 to 18 DIV. Imaging was done instantly in the warm Tyrode’s buffer.

### Immunostainings

For immunostainings (Figure 1), cells were fixed with 4 % (w/v) paraformaldehyde (PFA; P6148, Sigma-Aldrich) in 4 % Sucrose (0.12M) in PBS for 20 min at RT followed by washing three times with 1X PBS. Excess PFA was quenched with 100 mM NH_4_Cl for 20 min, then washed three times with PBS. The unspecific binding was blocked by incubating the coverslip in 2.5 % BSA and 0.1 % Triton X-100 in PBS for 15 min. The cells were then incubated in a humidified chamber at RT for 1 hour in primary antibodies diluted in 2.5 % BSA and 0.1 % Triton in PBS. Cells were washed twice with 2.5 % BSA and 0.1 % Triton X-100 in PBS and then, once with 2.5 % BSA in PBS. Finally, the coverslips were incubated for 1 hour in a humidified chamber at RT in secondary Abs (1:200, unconjugated from Jackson ImmunoResearch and conjugated in house as previously described (Coons et al., 1942), donkey anti-mouse IgG: cat# 715-005-151, RRID: AB_2340759; donkey anti-rabbit IgG: cat# 711-005-152, RRID: AB_2340585; donkey anti-guinea pig IgG: cat# 706-005-148, RRID:AB_2340443) in 2.5 % BSA in PBS. After washing the coverslips thrice with 2.5 % BSA in PBS, they were washed once with PBS containing high salt (400 mM NaCl), again washed with PBS, and finally mounted on slides with a mounting medium. Mounted coverslips were left at RT for a few hours, then kept at 4 °C overnight and imaged the following day. The primary antibodies used were anti-Syt1 clone 604.2 (105102, Synaptic Systems), anti-VGAT (Synaptic Systems, cat. #131103, RRID: AB_887870).

### General epi-fluorescence and confocal microscopy imaging

Images were acquired on Nikon Ti 2E (Nikon Corporation, Chiyoda, Tokyo, Japan) inverted epi-fluorescent microscope equipped with a 60X oil objective and Photometrics 95B camera. A dark Okolab (Ottaviano, Italy) cage incubator system was used to maintain 37 °C and 5 % CO2 (OKO Air Pump and CO2 controller). GFP/pHlourin was imaged in Alexa488 channel (EX 472/30, DM 495, BA 520/35), while mOrange2 probe was imaged in Cy3 channel (EX 531/40, DM 562 BA 593/40). Alexa647 was imaged in the Cy5 channel (EX 628/40, DM 660, BA 692/40). Imaging of fixed samples was also performed on a Zeiss laser scanning confocal microscope LSM710 with a scanning format of 1,024 x 1,024 pixels using a 63x oil-immersion objective with 1.3 numerical aperture (NA) at equal acquisition settings within each immunostaining.

### Stimulation assays for testing the effects of activity on the localization of SVs

Surface epitopes of Syt1 were saturated by incubating the neurons in divalent-free buffer (Tyrode buffer without Mg^2+^ and Ca^2+^) with HaloTag-free NbLumSyt1 for 5 min at 37 °C. Until fixation, neurons were kept at 37 °C on a heating metal plate. Before the stimulation, 15 µl of NbLumSyt1-HALO was premixed with 3 µl of Janelia Fluor 646 HaloTag ligand (100 pmol of JF646 stock at 33 µM, Promega GA112A) for 15 min at RT and diluted to 300 µl of final volume in Tyrode (to reach 4.85 µM final concentration of NbLumSyt1-HALO-JF646). Neurons were quickly washed with a drug buffer containing 50 µM APV and 10 µM CNQX to avoid recurrent activation. Neurons were stimulated at 20 Hz for 30 s. After stimulation, neurons were allowed to recover for 2 min in the same buffer, washed two times with Tyrode’s solution and fixed using 4 % PFA for 30 min at RT. The following steps were performed at RT. Neurons were quenched using 100 mM NH_4_Cl solution, quickly washed with PBS and blocked using blocking solution containing 2 % bovine serum albumin (BSA, PanReacAppliChem) and 0.1 % Triton X-100 (Sigma Aldrich, cat. 102604816) for 15 min. After, neurons were incubated in blocking solution containing mouse anti-bassoon antibodies (1:200 dilution, Enzo ADI-VAM-PS003-F, RRID: AB_1659574) for 1 hour. Then, neurons were washed three times in PBS for 5 min. After, neurons were incubated in blocking solution containing donkey anti-mouse Alexa Fluor 488 minimal cross-reactivity antibodies (AF488, 1:500 dilution, Jackson ImmunoResearch 715-545-151) and vGlut1-STAR580 nanobodies (1:500 dilution, FluoTag-X2 N1602-Ab580-L, NanoTag Biotechnologies, RRID: AB_3076007 (custom-labelling order)) for 1 hour. Then, neurons were washed three times in PBS for 5 min and mounted. The coverslips were dipped in ddH_2_O to remove salt excess, and the side of the coverslip was quickly dried on a Kim wipe tissue to remove excess liquid. Immediately after, the coverslips were mounted on a microscope slide using 10 µl of Prolong Glass Antifade mounting media (ThermoFisher, cat. P36980), left to harden overnight at RT and kept at 4 °C until imaging.

### Acquisition and analysis of images from stimulation assays

STED images were acquired using an Abberior microscope setup (Abberior Instruments GmbH) and Imspector software (version 16.3.14287-w2129). Using a UPLSAPO100XO objective (1.4 NA), 10 single z-plane images of 20 µm × 20 µm image size and 20 nm × 20 nm pixel size were collected per condition. AF488, STAR580 and JF646 fluorophores were excited using a 485 nm, 561 nm and 640 nm laser line and the emission was detected in the 500–550 nm, 605–625 nm and 650–720 nm range, respectively, in the STED mode. First, JF646 and STAR580 fluorophores were depleted using a 775 nm depletion laser line, immediately after AF488 fluorophores were depleted using a 595 nm depletion laser line. Image analysis was performed using an in-house written MATLAB script (The Mathworks Inc., v7.5.0.342 R2007b). Briefly, spots of NbLumSyt1 (recycling pool), vGlut1 (total vesicle pool) and Bassoon (active zone) were detected using pre-processing based on Fourier transformation. Bassoon was considered as a center of the detected spots, then NbLumSyt1 and vGlut1 spots were aligned with the Bassoon without modifying them separately. Finally, the detected spots were averaged and a 20 pixel (400 nm) thick intensity line profile vertically along the peak of the Bassoon was measured. An average image was calculated for every label, and a line profile was measured for each coverslip separately. Finally, raw fluorescence intensity profile values were median normalized for each coverslip and values measured in 5 pixels (100 nm) were pooled together and plotted as a boxplot. −100 to 300 nm distance range from the Bassoon peak was considered as proximal vesicle pool and 400 to 800 nm – as distal vesicle pool. Median values of the two pools were plotted separately for the 2 s, 30 s stimulation and the control condition. For visualization purposes in **Supplementary Fig. 2** an average image of all three coverslips was obtained.

### Image acquisition and analysis for “younger” and “older” vesicles

Confocal and STED images were acquired using an Abberior microscope setup (Abberior Instruments GmbH) including Olympus IX83 microscope body and Imspector software (version 16.3.15521-w2209). Using a UPLSAPO100XO objective (1.4 NA), 10 single z-plane images of 20 µm × 20 µm image size and 20 nm × 20 nm pixel size were collected per condition. Cy3 fluorophores were excited using a 561 nm laser line and detected in the confocal mode in the 605–625 nm range in the confocal mode. AF488 and JF646 fluorophores were excited with a 485 nm and 640 nm laser line and detected in the 500–550 nm and 650– 720 nm range, respectively, in the STED mode. Image analysis was performed using an in-house written MATLAB script (The Mathworks Inc., v7.5.0.342 R2007b). Briefly, “old” and “young” spots of NbLumSyt1 and active zone (Bassoon) spots were detected using pre-processing based on Fourier transformation. Bassoon was considered as a center of the detected spots, then “old” and “young” regions were aligned with the Bassoon without modifying them separately. Finally, the detected spots were averaged, and a 20 pixel (400 nm) thick intensity line profile was measured vertically along the peak of the Bassoon. An average image was acquired, and a line profile was measured for each coverslip separately. Finally, raw intensity profile values for each coverslip were median-normalized and values measured in 5 pixels (100 nm) were pooled together and plotted as a boxplot. For visualization purposes in the **Supplementary Fig. 2** an average image of all three coverslips was shown.

### Immunolabeling assay for Cntfr and lumenal domain of Syt1

To label Cntfr, 15 DIV primary hippocampal neurons were incubated with rabbit anti-CNTFRα (extracellular) antibodies (1:50 dilution, Alomone ACR-051, RRID: AB-2340925) in conditioned media at 37 °C for 40 min. Neurons were washed in cold Tyrode’s buffer and fixed using 4 % PFA for 30 min at room temperature (RT). The following steps were performed at RT. Neurons were quenched using 100 mM NH_4_Cl solution, quickly washed with PBS and blocked using blocking solution containing 2 % bovine serum albumin (BSA, Merck 1120180500) and 0.1 % Triton X-100 (TX-100, Merck 1122980101) for 15 min. After, neurons were incubated in blocking solution containing mouse anti-Syt1 luminal (1:200 dilution, Synaptic Systems, cat. #105201 604.1 clone, RRID: AB_2617068) antibodies for 1 hour. Then neurons were washed three times in PBS for 5 min. After, neurons were incubated in blocking solution containing anti-rabbit STAR635P (1:170 dilution, in-house conjugated rabbit Jackson ImmunoResearch 711-005-152 and STAR635P NHS ester Abberior 07679) and anti-mouse STAR580 (1:200 dilution, in-house conjugated mouse Jackson ImmunoResearch 715-005-151 and STAR580 NHS ester Abberior 38377) for 1 hour. Then neurons were washed three times in PBS for 5 min and mounted as described above.

Confocal and STED images were acquired using an Abberior microscope setup (Abberior Instruments GmbH) including Olympus IX83 microscope body and Imspector software (version 16.3.15521-w2209). Using a UPLSAPO100XO objective (1.4 NA), 10 single z-plane images of 20 µm × 20 µm image size and 20 nm × 20 nm pixel size were collected per condition. STAR580 and STAR635P fluorophores were excited using a 561 nm and 640 nm laser line, depleted using 775 nm laser line and the emission was detected in the 605–625 nm and 650–720 nm range, respectively, in the STED mode.

Image analysis was performed using in-house written ImageJ/Fiji (citation) macro. Briefly, binary masks of Syt1 and Cntfr positive regions were obtained by applying a 1 sigma radius Gaussian blur filter and setting a user-defined threshold. Such objects were filtered by area (0.04 – 2 µm^2^ for Syt1 and 0.01 – 0.8 µm^2^ for Cntfr) and counted for every image. An overlap between Syt1 and Cntfr was manually identified by selecting a fixed size circle ROI around the overlap. Such regions were counted and expressed as percentage over total Syt1.

### Proximity ligation assay (PLA) for Cntfr and Syt1

Labelling of Cntfr was performed as described above also including mouse anti-Syt1 luminal antibodies (1:150 dilution, Synaptic Systems, cat. #105 201, 604.1 clone, RRID: AB_2617068) in conditioned media at 37 °C for 40 min. In the control condition neurons were incubated with rabbit anti-ribeye (1:62.5 dilution, Synaptic Systems, cat. #192 003, RRID: AB_2261205) antibodies instead of Cntfr antibodies.

Neurons were washed, fixed, quenched and permeabilized as described above. After, neurons were subjected to proximity ligation assay using Duolink™ In Situ Orange Starterkit Mouse/Rabbit (DUO92102-1KT, Merck) according to the Duolink® PLA Fluorescence Protocol. After final washes, neurons were incubated in blocking solution containing guinea pig anti-Synaptophysin1 (1:500 dilution, Synaptic Systems, cat. #101 004, RRID: AB_1210382, polyclonal antiserum) antibodies for 1 hour. Then neurons were washed three times in PBS for 5 min. After, neurons were incubated in blocking solution containing anti-guinea pig AF488 (1:200 dilution, minimal cross-reactivity Jackson ImmunoResearch cat no: 706-545-148) antibodies for 1 hour. Then neurons were washed three times in PBS for 5 min and mounted as described above.

Epifluorescent images were acquired using an Olympus microscope setup including Olympus IX71 microscope body and OLYMPUS cellSens Dimension (version 2.3) software. Using a UPlanSApo 60× objective (1.35 NA), 10 single z-plane images of 147.92 µm × 111.59 µm image size and 0.1075 µm × 0.1075 µm pixel size were collected per condition. Fluorophores were excited using a Lumencor SOLA SE II lamp. Emission of Duolink Orange fluorophores were collected for 200 ms exposure time in a range with emission peak of 580 nm and emission of AF488 fluorophores were collected for 100 ms exposure time in a range with emission peak of 518 nm.

Image analysis was performed using in-house written ImageJ/Fiji macro. Briefly, binary masks of Synaptophysin and PLA positive regions were obtained by subtracting the background (rolling ball radius 10 pixels), applying a 1 sigma radius Gaussian blur filter on Synaptophysin signal and setting a user-defined threshold. Such objects were filtered by area (0.05 – 2 µm^2^ for Synaptophysin and larger than 0.03 µm^2^ for PLA) and counted for every image. An overlap between Synaptophysin and PLA was obtained by performing an “AND” operation between two masks and colocalizing regions were counted. Number of such regions was divided by the total number of Synaptophysin regions and normalized by the average of the control condition. Results were plotted as boxplots showing mean, 5th and 95th percentile of the dataset. Unpaired t test was performed using GraphPad Prism v9.4.0.673 for Windows (GraphPad Software, San Diego, California USA).

### Pharmacological alteration of Cntfr and Syt1 contacts

At 14 days in vitro (DIV) neurons were incubated either in conditioned medium only (control), or in conditioned medium supplemented with 8 nM ciliary neurotrophic factor (CNTF, C-245 Alomone Labs) for 2 hours or 24 hours, or 3 µM tetrodotoxin (TTX, 1069 Tocris) for 2 hours. As a proxy for neuronal activity, neurons were exposed to anti-Syt1 lumenal primary antibody directly labeled with ATTO647N (1:200, 105 311AT1 clone 604.2 Synaptic Systems) in conditioned medium for 30 mins at 37 °C. Later cells were washed, fixed, quenched, stained for Synaptophysin 1 and mounted as described above.

Imaging was performed using Plan Apo λ 60× oil (NA 1.4) objective and Andor DU-897 X-8536 camera of the inverted Nikon Ti microscope (Nikon Corporation, Chiyoda, Tokyo, Japan) setup. GFP and CY5 filters were used to excite the Alexa Fluor 488 and ATTO647N fluorophores accordingly. Images were collected of 135.21 × 135.21 µm size with a pixel size of 26.4 × 26.4nm using NIS Elements software.

All imaged were analyzed using a semi-automated Fiji macro. Synaptophysin positive regions were identified, and a common threshold was set. For general intensity analysis, Syt1 fluorescence intensity (a. u.) was measured in the whole synaptophysin positive region. For bouton specific analysis, Syt1 fluorescence intensity (a. u.) and area (µm^2^) was measured in every synaptophysin positive bouton. 2 µm^2^ maximum area filter was applied to exclude boutons that might have been merged during the thresholding. Boutons were divided into three groups: small (<0.4 µm^2^), medium (0.4-0.9 µm^2^) and large (>0.9 µm^2^). An average of the mean intensity was calculated for every group in every image (n = 30 images), in every condition (N = 3 coverslips), normalized to the median of the control and expressed as percentage. One-way ANOVA followed by post hoc Bonferroni multiple comparisons test was performed using GraphPad Prism v9.4.0.673 for Windows (GraphPad Software, San Diego, California USA).

### Electrical stimulation of cultured neurons

A perfusion-type electrical stimulation magnetic chamber (EC B18, Live Cell Instrument, Gyeonggi-do, The Republic of Korea) was used to achieve field stimulation at indicated frequencies and pulses with A310 Accupulser Stimulator (Precision Instruments, Sarasota, FL, USA). Stimulation was performed in the presence of CNQX and APV to reduce recurrent activity from excitatory synapses. 1 image every 2 seconds was captured for a total of 3.5 min. After the first 30 seconds, electrical stimulation was applied and 3 min later, 50 mM NH_4_Cl was applied. For the Bafilomycin A1 control experiment, 250 nM Bafilomycin A1 was included right from the beginning of the image acquisition, in the Tyrode’s buffer. Exposure times and light intensity were adjusted to obtain minimum photobleaching. ‘Perfect Focus System’ feature was used before the start of the experiment, to avoid any focal drift caused due to stimulation or addition of NH_4_C or change of pH/Cl-clamped buffers.

### Electrophysiology

Whole-cell recordings of pyramidal neurons were achieved using Axopatch 200B and with the Clampex 8.0 software (Molecular Devices), filtering at 1 kHz and sampling at 5 kHz in voltage-clamping at −70 mV. Only experiments with access resistance values of 5–20 MΩ were considered for analysis. All recordings were performed at room temperature. The composition of the internal pipette solution was: 115 mM CsMeSO_3_, 20 mM tetraethylammonium chloride, 10 mM CsCl, 10 mM HEPES, 5 mM NaCl, 0.6 mM EGTA, 4 mM Mg-ATP, 0.3 mM Na2GTP and 10 mM QX-314 (lidocaine N-ethyl bromide). The final solution was adjusted to a pH of 7.3 and an osmolarity of 305-310 mOsM. The final resistance of the electrode tips used was ∼2-5 MΩ. To elicit evoked responses, electrical stimulation was delivered using a parallel bipolar electrode (FHC) and a constant current unit (WPI A385) coupled with a Master-8 controller (A.M.P.I.). The pulse duration was 0.1 ms and the intensity was 35 mA (the same stimulation equipment and parameters were used for the live fluorescence imaging and electrophysiology experiments in **Fig. 3a-g**). The extracellular solution was a modified Tyrode’s solution containing 150 mM NaCl, 4 mM KCl, 10 mM glucose, 10 mM HEPES and 2 mM MgCl2, adjusted to pH 7.4 and 315-320 mOsM. The agonists against ionotropic glutamate receptors 6-cyano-7-nitroquinoxaline-2,3-dione (CNQX, 10 μM) and aminophosphonopentanoic acid (AP-5, 50 μM) were used to isolate inhibitory postsynaptic currents (IPSC). To isolate excitatory (AMPA-mediated) postsynaptic currents (EPSC), the GABA-A receptor inhibitor picrotoxin (PTX, 50 μM) and AP-5 were added to the bath solution. Spontaneous neurotransmission was recorded with the addition of 1 μM TTX. Evoked recordings were analyzed using Clapfit (Molecular Devices) and miniature events were analyzed using MiniAnalysis (Synaptosoft).

### Nanobody uptake and post hoc immunostainings

For NbLumSyt1 live cell uptake at DIV 14, the corresponding coverslip was incubated in 200 µL conditioned medium diluted 1:200 with GFP-tagged nanobody for 30 min at 37 °C and 5 % CO2. After incubation, the coverslip was transferred into the original medium for 15 min and for labeling, neurons were three times washed with 1x PBS and fixed with 4 % PFA/ 4 % sucrose in PBS for 15 min at room temperature (RT). Each coverslip was washed three times with PBS, blocked with blocking solution (1x PBS, 10 % normal goat serum (NGS) and 0.3 % Triton X-100) for 30 min at RT and incubated with primary antibodies (anti-GFP (rabbit polyclonal, used at 1:1000 in IF, Abcam, Cat# ab6556, RRID:AB_305564), anti vGLUT1 (guinea pig polyclonal, used at 1:500 in IF, Synaptic Systems, Cat# 135 304, RRID:AB_887878)) diluted in blocking solution for 2 h at RT. Before and after incubation with the corresponding secondary antibodies (goat anti-rabbit IgG Alexa Fluor 488 (polyclonal, used at 1:400, Thermo Fisher Scientific, Cat# A-11008, RRID:AB_143165), goat anti-guinea pig IgG Alexa Fluor 568 (polyclonal, used at 1:400, Thermo Fisher Scientific, Cat# A-11075, RRID:AB_141954)) for 45 min, each coverslip was washed three times with PBS and finally mounted on glass slides in Immumount (Thermo Fisher Scientific).

### Automated solid-phase peptide synthesis

The μSPOT peptide arrays ^65^ were synthesized using a MultiPep RSi robot (CEM GmbH, Kamp-Lintford, Germany). As a substrate we used in-house produced, acid labile, amino-functionalized, cellulose membrane discs containing 9-fluorenylmethyloxycarbonyl-β-alanine (Fmoc-β-Ala) linkers (with an average loading of 130 nmol/disc with spots of 4 mm in diameter). Synthesis was initiated by Fmoc deprotection using 20% piperidine (pip) in dimethylformamide (DMF) followed by washing with DMF and ethanol (EtOH). Peptide chain elongation was achieved using a coupling solution consisting of preactivated amino acids (aa, 0.5 M) with ethyl 2-cyano-2 (hydroxyimino) acetate (oxyma, 1 M) and N,N’-diisopropylcarbodiimide (DIC, 1 M) in DMF (1:1:1, aa:oxyma:DIC). Couplings were carried for 3 × 30 min, followed by capping (4% acetic anhydride in DMF) and washes with DMF and EtOH. Synthesis was finalized by deprotection with 20% pip in DMF (2 × 4 μL/disc for 10 min each), followed by washing with DMF and EtOH. Dried discs were transferred to 96 deep-well blocks and treated, while shaking, with side-chain deprotection solution, consisting of 90% trifluoracetic acid (TFA), 2% dichloromethane (DCM), 5% H_2_O and 3% triisopropylsilane (TIPS) (150 μL/well) for 1.5 h at room temperature (rt). Afterward, the deprotection solution was removed, and the discs were solubilized overnight (ON) at rt, while shaking, using a solvation mixture containing 88.5% TFA, 4% trifluoromethanesulfonic acid (TFMSA), 5% H_2_O and 2.5% TIPS (250 μL/well). The resulting peptide-cellulose conjugates (PCCs) were precipitated with ice-cold ether (0.7 mL/well) and spun down at 2000 g for 10 min at 4 °C, followed by two additional washes of the formed pellet with ice-cold ether. The resulting pellets were dissolved in DMSO (250 μL/well) to provide the final stocks. PCC solutions were mixed 2:1 with saline–sodium citrate (SSC) buffer (150 mM NaCl, 15 mM trisodium citrate, pH 7.0) and transferred to a 384-well plate. For transfer of the PCC solutions to white-coated CelluSpot blank slides (76 × 26 mm, Intavis AG Peptide Services GmbH and CO. KG), a SlideSpotter (CEM GmbH) was used. After completion of the printing procedure, slides were left to dry ON.

### Peptide microarray binding assay

The microarray slides before use were blocked for 60 min in 5% (w/v) skimmed milk powder (Carl Roth) in phosphate-buffered saline (PBS; 137 mM NaCl, 2.7 mM KCl, 10 mM Na_2_HPO_4_, 1.8 mM KH_2_PO_4_, pH 7.4). After blocking, slides were incubated for 30 min with 7.36 nM of ALFA-NbLumSyt1 in blocking buffer, then washed three times with PBS. The ALFA-NbLumSyt1 was detected with a secondary 1:50000 diluted nanobody anti-ALFA-HRP (NanoTag Biotechnologies, cat. N1505-HRP, RRID: AB_3075989). The nanobodies were applied in a blocking buffer for 30 min, with three PBS washes between the nanobodies and the application of the secondary nanobody. The chemiluminescent readout was obtained using SuperSignal West Femto maximum sensitive substrate (Thermo Scientific GmbH, Schwerte, Germany) with a c400 Azure imaging system (lowest sensitivity, 10 s exposure time). Binding intensities were quantified with FIJI ^66^ using the “microarray profile” plugin (OptiNav Inc, Bellevue, WA, USA). The raw grayscale intensities for each position were obtained for the left and right sides of the internal duplicate on each microarray slide, n = 4 arrays in total.

### Expression and purification of the nanobody-Syt1 luminal peptide complex

The DNA sequences encoding NbLumSyt1and 6x histidine-tagged Syt1 luminal peptide (MGEGKEDAFSKLKEKFMNELHKGHHHHHH) were cloned into a pETDuet-1 vector. The NbLumSyt1-Syt1 luminal peptide complex was expressed in *E. coli* BL21(DE3) using standard IPTG induction and purified by Immobilized metal affinity chromatography (IMAC) followed by Size exclusion chromatography (SEC) using a Superdex75 16/60 column (GE Healthcare) equilibrated with 20 mM HEPES pH 7.4, 150 mM NaCl, 5 % (v/v) glycerol. The purified complex was concentrated to 43 mg/ml for crystallization.

### Crystallization and structure of the nanobody-Syt1 luminal peptide complex

The NbLumSyt1 - luminal peptide complex (43 mg/mL) was crystallized at 21 °C via vapor diffusion in a 24-well sitting drop plate, where 1µL of protein solution was mixed with 1µL of reservoir solution (1.6 M DL-malic acid). X-ray diffraction data was collected at station I04 of the Diamond Light Source (Oxford, UK) equipped with a Dectris Eiger2 XE 16M detector. The data was processed and scaled with DIALS ^67^ and Aimless ^68^ within the CCP4 suite ^69^. Molecular replacement was performed in Phaser ^70^ using the structure with PDB code 6I2G as the search model ^23^. Several cycles of model building and refinement were carried out using Coot ^71^ and Refmac5 ^72^. Data collection and processing statistics are presented in **Supplementary Table 2**. Figures and analysis are based on chains A (nanobody) and I (peptide) which have the best electron density in the structure. The coordinates and structure factors for NbLumSyt1 were deposited in the PDB under the accession code 8B8I.

### Coupling of the NbALFA with nanogold particles

As described above, NbALFA was produced in SHuffle cells, bearing a free ectopic Cys on its C-terminus. NbALFA-Cys was reduced using 10 mM TCEP for 1h. The excess of TCEP was removed using a Nap5 (Cytiva) column and immediately mixed with an equimolar amount of monofunctionalized maleimide 3 nm gold particles (Nanopartz, Loveland, US) and left to react for two hours at room temperature. Non-conjugated gold was removed using a size exclusion column (Superdex 75 increase 10/300 GL, Cytiva). NbALFA-Gold was kept at 4 °C.

### High-pressure freezing, freeze substitution, and ultramicrotomy

Sapphire discs with neurons were incubated in pre-mixed NbLumSyt1-ALFA-tag Nb:NbALFA-Gold complex in a conditioned medium or in conditioned medium containing only NbALFA-Gold for 45 min. The neurons then underwent washing steps in pre-warmed (37 °C) conditioned medium before rapid cryofixation using EM ICE high-pressure freezer (Leica) and storage in liquid nitrogen. Semi-automated freeze-substitution was performed using an AFS2 device (Leica) equipped with customized aluminum sapphire disc revolvers and samples were subsequently embedded in epoxy resin according to a previously published protocol ^73^. One minor modification implemented to enhance membrane contrast entailed the addition of 1% water ^74^ to the sequential freeze-substitution cocktails of 0.1% tannic acid and 2% osmium tetroxide in glass-distilled acetone. Upon sapphire disc removal from polymerized samples, block-faces were trimmed with an EM TRIM high-speed milling device (Leica) in preparation for ultramicrotomy. A 35° Ultra Semi diamond knife (Diatome) mounted on a UC7 ultramicrotome (Leica) was used to cut alternating series of sections onto formvar-coated copper mesh grids at two different thicknesses: (i) 60 nm-thick sections were visually inspected to assess ultrastructural preservation and synapse density under an 80kV LEO 912 transmission electron microscope (Zeiss), and (ii) 250 nm-thick sections were coated on both surfaces with Protein A-conjugated 10 nm gold fiducial particles (Cell Microscopy Core Products, University Medical Center Utrecht, The Netherlands) in preparation for transmission electron tomography.

### Electron microscopy

Synapses were identified and selected for 3D ultrastructural analysis in tiled 11000 x magnification overviews acquired using a 200 kV Talos F200C G2 scanning/transmission electron microscope equipped with a 16 MP Ceta CMOS camera (Thermo Scientific) and MAPS v3.1 software (Thermo Scientific). Tilt series (±60° tilt range; 1° increments) were acquired at 36000 x (unbinned pixel size = 0.4 nm) or 57000 x magnification (unbinned pixel size = 0.25 nm) from orthogonal axes using a Model 2040 dual-axis specimen holder (Fischione) and TEM Tomography v4 software (Thermos Scientific). Tomograms were generated from tilt-series using gaussian filtering and a weighted back-projection algorithm implemented using *Etomo* software IMOD software package (Kremer *et al.,* 1996). The acquired 36000 x magnification (n=39 and n=36 for NbALFA-Gold and NbLumSyt1-ALFA-tag:NbALFA-Gold complex, respectively) and 57000 x magnification tomograms (n=11 for NbLumSyt1-ALFA-tag:NbALFA-Gold complex) were analyzed using 3dmod software and additional programs within the IMOD software package ^75^. For tomograms acquired at 36000 x magnification, gold particles were detected using *findbeads3d* and were sorted based on their location (intracellular, extracellular, presynaptic, postsynaptic, SVs, endosomes and multivesicular bodies). For tomograms acquired at 57000 x magnification, the volumes of individual and clustered gold particles were quantified using *contourmod* and *imodinfo*. Active zones and clear-core vesicles were segmented using *3dmod*, and their diameters and closest approach distances were quantified using *imodinfo* and *mtk* programs, respectively.

### Live imaging of synaptic vesicle exo- and endocytosis

For initial experiments, fluorescence was recorded using a Nikon Eclipse TE2000-U microscope (Nikon) and an Andor iXon+ back-illuminated EMCCD camera (Model no. DU-897E-CSO-#BV). For illumination, we used a Lambda-DG4 illumination system (Sutter instruments) with a FITC emission filter. Images were acquired at 5 Hz. Solutions were perfused using an automatic, constant flux system (AutoMate Scientific). Circular regions of interest (ROI) of 2 µm diameter were drawn around local fluorescence maxima (putative presynaptic boutons) and measured using Fiji (NIH). Fluorescence peaks synchronous respect to the stimulation were detected and analyzed using Matlab. For additional experiments, live and actively recycling neurons were continuously incubated for 15-30 min or transfected with Syt1-pHluorin. Neurons were then stimulated with 200 action potentials (APs) delivered at 10 Hz or 40 Hz frequencies. Fluorescence intensity of pHluorin was monitored over time using live imaging to track SV exo- and endocytosis. For experiments comparing the kinetics of SV exo- and endocytosis, the average time course of the Syt1 response was recorded following stimulation with 200 APs at 10 Hz. Fluorescence changes were normalized to the stimulation peak, and comparisons were made between neurons stained with the Syt1 pHluorin Nb and those overexpressing Syt1 pHluorin. No differences in the kinetics were observed between the two conditions at this stimulation frequency. Fluorescence recovery to baseline was quantified at 120 seconds post-stimulation from the fluorescence transients obtained. The average time course of the Syt1 response after stimulation with 200 APs at 40 Hz was recorded and normalized to the stimulation peak. To assess the relative amount of SV exocytosis, fluorescence changes were normalized to the total Syt1 pool, revealed by the application of NH_4_Cl, following stimulation with 200 APs at 10 Hz. Surface expression of Syt1 pHluorin was assessed by measuring the fraction of Syt1 pHluorin on the cell surface, with fluorescence normalized to 1 in NH_4_Cl buffer and 0 in acidic buffer. Fluorescence data were presented as means ± standard error of the mean (SEM). N values represent the number of individual coverslips from three separate neuronal preparations, as indicated in the plots. Statistical significance in these experiments was determined using a two-tailed Student’s t-test, with significance thresholds set at ns P > 0.05, ∗P < 0.05.

For additional experiments live and actively recycling neurons were always incubated for 15-30 min or transfected with Syt1 pHluorin. Fluorescence intensity of pHluorin was monitored over time using live imaging to track SV exo- and endocytosis. The cultures were continuously perfused with imaging buffer (mM concentrations: 136 NaCl, 2.5 KCl, 2 CaCl_2_, 1.3 MgCl_2_, 10 glucose, 10 HEPES, pH 7.4). Imaging was performed on an inverted Zeiss Axio Observer Z1 microscope equipped with a 40x EC Plan-Neofluar oil immersion objective (NA 1.3), Colibri 7 LED light source, and AxioCam 506 camera, controlled by ZEISS ZEN 2 software. Transfected neurons were imaged at 2 Hz using a GFP filter (excitation: 450–490 nm; beam splitter: 495 nm; emission: 500–550 nm). For experiments comparing the kinetics of SV exo- and endocytosis, the average time course of the Syt1 response was recorded following stimulation with 200 action potentials (AP) at either 10 Hz or 40 Hz. Fluorescence changes were normalized to the stimulation peak, and comparisons were made between neurons stained with the Syt1 pHluorin Nb and those overexpressing Syt1 pHluorin. Fluorescence recovery to baseline was quantified at 120 seconds post-stimulation from the fluorescence transients obtained. To assess the relative amount of SV exocytosis, fluorescence changes were normalized to the total Syt1 pool, revealed by application of NH_4_Cl (50mM), following stimulation. Surface expression of Syt1 pHluorin was assessed by measuring the fraction of Syt1 pHluorin on the cell surface, with fluorescence normalized to 1 in NH_4_Cl buffer and 0 in acidic buffer (substituting 20 mM MES for HEPES, pH 5.5). Fluorescence data were processed offline using the Time Series Analyzer plugin for ImageJ (https://imagej.nih.gov/ij/plugins/time-series.html) and presented as means ± standard error of the mean (SEM). N values represent the number of individual coverslips from three separate neuronal preparations, as indicated in the plots. Statistical significance in these experiments was determined using a two-tailed Student’s t-test, with significance thresholds set at ns P > 0.05, ∗P < 0.05.

### In vivo pH calibration and resting pH estimation

The in vivo pH calibration profile and resting pH were estimated according to ^28^ with some modifications. In addition, under the assumption that the H^+^ and Cl^−^ ions can move freely when the membranes are perfused with blockers and ionophores, the pH^−^ concentration inside the probe containing SV should be equal to the external bath solution. The current technique has been used in virtually all previous pH studies in the literature ^28,76,77^.

For calibrating the internalized pH-sensitive probes, neurons were treated with an ionophore cocktail containing the proton uncoupler FCCP, the K^+^ ionophore valinomycin, the V-ATPase inhibitor bafilomycin A1, the membrane solubilizer Triton X-100, and the K^+^/H^+^ exchanger nigericin to perfuse the membranes in high K^+^ (122.4 mM) buffers adjusted to a defined pH. Images acquired at each pH were analyzed by bouton analysis. A pH-calibration profile was then plotted to obtain the pKa of each pH probe.

For estimating the intravesicular pH, neurons preloaded with the pHluorin/mOrange 2 probes were treated with the TEV-protease for 3 min to remove all external fluorescence signal outside SVs, which was defined as the “resting state”. They were then very briefly incubated in ionophores containing two pH-adjusted buffers - maximum and minimum pH as based on the pH calibration curve. The images were captured immediately and were referred to as ‘max’ and ‘min’. Furthermore, cells in the same region of interest (ROI) were immunostained with primary labeled Abs against vGlut and vGAT to differentiate the synaptic populations into Glutamatergic and GABAergic. Anti-vGLUT1 (guinea pig polyclonal, used at 1:500 in IF, Synaptic Systems, cat. #135304, RRID: AB_887878) anti-VGAT (Synaptic Systems, cat. #131103, RRID: AB_887870, luminal domain). The ROI was preserved by performing the experiment on top of the microscope stage and saving the X-Y position for every image that was captured.

These images were analyzed and fitted with the following equation:

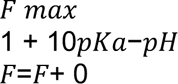

The pKa was calculated from the pH calibration data. F is the fluorescence readout at ‘resting’ condition, whose pH is unknown. F0 and Fmax are the fluorescence values at the minimum and maximum pH, respectively, whose pH is known; thus, the unknown resting pH is calculated.

### APEX2 labelling for mass spectrometry

APEX2 labeling was performed with minor modifications as previously described ^29^. As a negative control a NbALFA-APEX2, which did not recognize any tag in our WT neurons, was used. Biotin-phenol was prepared by dissolving 50 mg BP (Sigma SML2135-50MG) in 276 µl DMSO, followed by sonication in a water bath for 20 min to complete dissolution. This yielded 500 mM BP stock, which was further diluted in 10 µl aliquots and stored at −80 °C until use. For H_2_O_2_ labeling, a fresh 100X stock was prepared for each use by adding 2 µl of a 30% H_2_O_2_ solution to 192 µl of Tyrode’s solution to obtain a 100 mM stock. This stock was diluted 1:100 in the labeling reaction to a final concentration of 1 mM. The quencher solution consisted of 10 mM sodium ascorbate, 5 mM Trolox, and 10 mM sodium azide in Tyrode’s solution. Sodium ascorbate was freshly dissolved in Millipore water to a concentration of 1 M, while Trolox was similarly prepared at 500 mM and sonicated. A 1 M stock solution of sodium azide was prepared, and aliquots were frozen at −20°C. To prepare the quencher solution, 19.4 ml of Tyrode’s was added to 200 µl of freshly prepared sodium ascorbate and Trolox and frozen sodium azide stock. This solution was prepared fresh as they are only effective for a short period. For these experiments, primary neuron cultures were prepared in 10 cm plates at a concentration of 5 M per plate and treated with a BP labeling protocol. Media was removed from each plate and stored at 37°C, leaving 1 ml of media on the cells. Three labeling cocktails were prepared with BP at a 1:500 dilution in pre-conditioned media in the presence of Nb-APEX2, Nb control or without nanobody addition. These were then drop-labeled onto neurons. After incubation for 30 min in an incubator with gentle shaking, the neurons were washed three times with their original medium and then with BP buffer (20 ml Tyrode with 20 µl BP). A fresh H_2_O_2_ pulse solution was prepared by adding 80 µl of the 100X H_2_O_2_ stock to 8 mL of BP buffer, and 2 mL of this solution was applied to each Petri dish. After incubation for 30 seconds, the H_2_O_2_ solution was removed, and the reaction was quenched by adding 2 ml of the prepared quencher solution. Later, the previous quencher solution was replaced with 1 ml for the effect of cell scraping. The cells collected in the quencher solution were pelleted by centrifugation at 2000 rpm for 2 min, and the supernatant was discarded. The resulting cell pellets were snap-frozen and stored at −80 °C for subsequent lysis and bead enrichment procedures.

### Bead enrichment of biotinylated proteins

For cell lysis, 20 mL of RIPA lysis buffer was freshly prepared with Millipore water. The buffer contained 50 mM Tris, 150 mM NaCl, 0.1% SDS (wt/vol), 0.5% sodium deoxycholate (wt/vol), 1% Triton X-100 (vol/vol), a protease inhibitor cocktail, 1 mM PMSF, 5 mM Trolox, 10 mM sodium azide, and 10 mM sodium ascorbate. The pH was then adjusted to 7.5 with HCl and the buffer was stored at 4 °C. Prior to lysis, the centrifuge was pre-cooled to 4°C and liquid nitrogen and ice were provided. Cells were lysed in freshly prepared RIPA lysis buffer, using ∼300-400 µl per cell pellet. Lysates were gently pipetted and kept on ice or at 4°C throughout the process to preserve protein integrity. After 5 min of incubation on ice, the streptavidin magnetic bead slurry was washed with 1 mL of RIPA lysis buffer for each of two washes per 50 µL used per experimental set. Lysates were then cleared by centrifugation at 15.000 g for 10 min at 4°C. A fraction of the total homogenate (50-80 µl) was removed and snap-frozen for later MS processing and analysis. The remaining 300 µl of the sample was incubated with the washed beads for 2 hours at RT with rotation. After incubation, the beads were pelleted using the magnetic rack, and the supernatant was retained and snap-frozen for future use, if required. The beads were then thoroughly washed with 1 ml RIPA Lysis Buffer at 4 °C. The beads were completely resuspended, spun on a magnetic rack and pelleted. After washing, all wash buffer was removed before the next wash. This was repeated twice with RIPA Lysis Buffer, transferring to a new tube for each resuspension. Subsequent washes were performed with 1 ml of 1 M KCl, 1 ml of 0.1 M Na_2_CO_3_, 1 ml of 2 M urea in 10 mM Tris-HCl at pH 8.0, and two additional washes with 1 ml of RIPA lysis buffer. All wash buffers were kept on ice during the washing steps to ensure sample stability.

### EM confirmation of NbLumSyt1-APEX2 localization

Following the biotinylation step, the samples were washed thoroughly with PBS to remove any unreacted biotin-phenol and residual quenching agents. This washing step was repeated multiple times to ensure the complete removal of any non-specifically bound substances. The samples were then incubated with a solution of streptavidin conjugated to horseradish peroxidase (HRP) at a concentration of 1 μg/mL for one hour at room temperature with gentle agitation. This incubation step allowed the streptavidin-HRP complex to bind specifically and efficiently to the biotinylated proteins, facilitating subsequent detection steps. The cells were immobilized with 2.5 % glutaraldehyde in 0.1 M cacodylic buffer (pH 7.4, Electron Microscopy Sciences) for one hour at room temperature. Fixation was completed at 4 °C overnight. Subsequently, the cells were washed three times with ddH2O to remove fixative followed by incubation with DAB (1 mg/ml, Electron Microscopy Sciences) in ddH2O for 5 min on ice. Next, the supernatant was discarded and the cells incubated in an ice-cold solution of DAB (1 mg/ml) with H_2_O_2_ (final concentration of 5.88 mM) in ddH2O for 30 min on ice. For post-fixation and staining, the cells were incubated in 1 % OsO4 for about 2 min on ice. Subsequently, the cells were washed several times with ddH2O and dehydrated with a graded ethanol series (30, 50, 70, 100 %) with two final dehydration steps in propylene oxide for 5 min each. Finally, the cells were infiltrated with EMBed-812 resin (Science Services) and cured at 60 °C for 48 hours. TEM micrographs of thin sections (60-70 nm thickness) were recorded on a Phillips CM120 TEM microscope equipped with a 2k x 2k slow scan CCD camera (TVIPS) and operated at 120 kV.

### LC-MS/MS analysis

MS analysis of digested proteins was performed in three biological replicates, each analyzed in technical duplicates, using a hybrid quadrupole-ion trap-orbitrap mass spectrometer (Orbitrap Fusion, Thermo Fisher Scientific, San Jose, USA). Data acquisition was performed with a data-dependent acquisition method. Peptides were first concentrated on a C18 PepMap100 trapping column (0.3 mm × 5 mm, 5 μm, Thermo Fisher Scientific, Waltham, USA) and subsequently separated on an in-house packed C18 analytical column (75 μm × 300 mm, Reprosil-Pur 120 C18-AQ, 1.9 μm, Dr. Maisch GmbH, Ammerbuch, Germany). Liquid chromatography was performed on an UltiMate 3000 UHPLC nanosystem (Thermo Fisher Scientific, Waltham, USA) with columns pre-equilibrated in a mixture of 95% buffer A (0.1% v/v formic acid in water) and 5% buffer B (80% v/v acetonitrile with 0.1% v/v formic acid in water). Peptides were eluted over a 118-minute gradient from 5 to 50% buffer B, followed by a 90% buffer B wash for 6 min and re-equilibration at 5% buffer B for another 6 min. The mass spectrometer was operated in DDA mode with MS1 scans covering a 350-1650 m/z range, at a resolution of 120,000 at 200 m/z, with a 300% automated gain control (AGC) target, and a maximum injection time of 50 ms. Peptide ions with charge states of 2-7 were selected for MS/MS analysis using a 1.6 m/z isolation window. Fragmentation was induced by HCD at 28% of the normalized collision energy (NCE); fragments were detected in the Orbitrap at a resolution of 15,000 at 200 m/z, with an AGC of 1000% normalized and a maximum injection time of 54 ms. The duty cycle was maintained at 2.5 seconds with dynamic exclusion of 30 seconds. Subsequently, the analysis of the raw data was performed in MaxQuant using the *Rattus norvegicus* proteome (UP000234681), supplemented by sequences of the APEX2 nanobodies as the positive controls, in the validation of IP. For statistical treatment of the data the label-free quantification (LFQ) intensities of the proteins found significantly enriched vs. both input and the negative controls IP (an NbALFA-APEX2 IP or a no-Nanobody IP) were selected. For initial analyses we relied on unbiased gene ontology cellular component analysis ^78^. Since several potential interactors were found, we decided to concentrate on the targets that were strongly enriched both vs. the controls (>1500 enrichment vs the controls) and enriched even when compared to the initial input (>1.5 enrichment vs the input). Only few proteins passed this stringent cutoff, including Cntfr, the immunoglobulin superfamily member 8 (Igsf8), the integral membrane protein 2B (Itm2b) and the heat shock 70 kDa protein 13 (Hspa13).

### Pre-complex formation of nbSyt1 nanobodies for uPAINT imaging

The NbLumSyt1-pH/Nb-Atto647N and NbLumSyt1-HALO/JF549 complexes were freshly prepared before each experiment. To prepare the NbLumSyt1-pH/nb-At647N complex, a 110-time molar excess of NbLumSyt1-pH was mixed with anti-GFP NbLumSyt1-At647N (Camelid sdAB from Synaptic Systems, N0301-At647N-S) at 4°C for 16 hours, covered from light, in gentle rotation (20 rpm). The NbLumSyt1-pH/nb-At647N complex was then centrifuged at 18,000g for 10 min at room temperature to remove precipitates, and subsequently diluted (1:200) in High K^+^ buffer, mixed at room temperature for 5 min in gentle rotation (20 rpm), and used for imaging on the same day. To prepare the NbLumSyt1-HALO/JF549 complex, a 110-time molar excess of NbLumSyt1-HALO was mixed with JaneliaFluor 549 HaloTag ligand for super-resolution microscopy (Promega, GA1110) at room temperature (i.e., 22-25°C) for 30 min, covered from light, in gentle rotation (20 rpm). The NbLumSyt1-HALO/JF549 complex was then centrifuged at 18.000 g for 10 min at RT to remove precipitates, and subsequently diluted (1:200) in High K^+^ buffer, mixed at RT for 5 min in gentle rotation (20 rpm), and used for imaging on the same day. These approaches ensure a 1 nM effective concentration of the fluorescently labeled nanobody (NbLumSyt1-pH/Nb-At647N or NbLumSyt1-HALO/JF549) on the imaging plate, allowing partial labeling of the Syt1 nanobodies without over-saturating the fluorescent signal enabling single molecule detection.

### Single molecule imaging by uPAINT

The single molecule uPAINT experiments to detect and track endogenous and over-expressed Syt1 on the plasma membrane were performed as described previously ^34,79^. Time-lapse imaging was carried out at 50 Hz and 20 ms exposure at 37 °C on Roper microscope (Roper Scientific) equipped with an iLas2 double-laser illuminator, a CFI Apo TIRF 100× 1.49 NA objective (Nikon) and an Evolve512 delta EMCCD camera (Photometrics, model no. Evolve 512 Delta). A quadruple beam splitter (LF 405/488/561/635-A-000-ZHE; Semrock) and a QUAD band emitter (FF01-446/510/581/703-25; Semrock) were used, and image acquisition was performed using MetaMorph Microscopy Automation and Image Analysis Software (v7.7.8, Molecular Devices). The single-molecule localization and dynamics were extracted from 16,000 frames, acquired in HILO as previously described ^80^. Endogenous Syt1-bound NbSyt1-pH/Nb-At647N and nbSyt1-HALO/JF549 complexes and over-expressed Syt1-pH-bound nb-At647N were detected and tracked using a combination of wavelet segmentation ^81^ and optimization of multi-frame object correspondence by simulated annealing ^82^. Single molecule localization and tracking was done in PALMTracer ^83^ software within the MetaMorph Software. We applied a cross-correlation-based drift correction to the data using SharpViSu tool ^84^, and tracks shorter than seven frames were excluded from the analysis to minimize nonspecific background. The colour coding for the super-resolved images was performed using Fiji/ImageJ ^66^ (2.0.0-rc-68/1.52h; National Institutes of Health). Diffusion coefficients were calculated for each trajectory and presented in a color-coded pixel at the site of localization. In the average intensity maps, each pixel indicates the localization of an individual detected molecule. The area with the highest density is represented in white. The color coding of the trajectory maps is arbitrary and corresponds to the colors shown in the graphs. Statistics for these experiments considered normality and lognormality, as well as AUC and frequency distribution quantifications. These tests were carried out in Prism 8 for macOS Software (version 8.3.0). Statistical analysis of normally distributed data was performed using ONE-way ANOVA Multiple comparison, and for non-normally distributed data, ONE-way ANOVA Kruskal-Wallis test Multiple comparison. Error bars are ± SEM for independent experiments and individual dots in the scatter plot graphs represent individual neurons. N = 22 neuronal cultures from 3 independent experiments in each condition. For DBSCAN, 10 representative datasets from nbSyt1-HALO/JF549 experiments were analysed. Non-significance is indicated with n.s., and asterisks indicate the following p-values *p < 0.05, ** p < 0.01, and **** p <0.0001. N = 22 neuronal cultures from 3 independent experiments in each condition.

### Nanocluster analysis

Nanoclustering analysis of Syt1 uPAINT data was performed using the Python based Nanoscale Spatio-Temporal Indexing Clustering (NASTIC) workflow ^37^. NASTIC determines clustering based on overlapping spatiotemporal bounding boxes of each tracked single molecule in space and time. Briefly, trajectories within a region of interest (ROI) were screened to remove those with less than 8 sequential detections. Spatial centroids were determined for each trajectory by averaging the x and y co-ordinates of all the detections. A convex hull of all the detections associated with the trajectory was calculated, and the radius R was derived from its area. An idealized 3D spatiotemporal bounding box with [x,y,z] dimensions x = 2R*r*, y = 2R*r* and z = *t* was created around each centroid, where *r* = 1.2 and *t* = 20 s. These *r* and *t* values have been previously empirically determined to best yield clustering metrics reflecting the ground truth of a range of synthetic trajectory datasets, as well as experimentally derived single-molecule imaging data. Bounding boxes were indexed into a 3D R-tree spatial database, which was then queried to return overlapping bounding boxes. Overlapping bounding boxes were then used to assign their parent trajectories into clusters. For each NASTIC cluster, a convex hull of all the detections comprising the clustered trajectories was used to determine the cluster area, radius and subsequent metrics. Clusters with radii > 150 nm were screened from the analyses.

### MoNaLISA imaging scheme and acquisition

The MoNALISA setup used in this study was custom-built, as previously reported ^40^. Briefly, images have been recorded with a multifoci pattern of periodicity 625 nm coupled with an OFF pattern of 312.5 nm. The ON-switch was performed with 405 nm light 650 W/cm^2^ for 0.5 ms, the OFF confinement with 4 ms of 488 nm light at 650 W/cm^2^ and finally the “read out” with 240 kW/cm^2^ of 488 nm per 1 ms. The step size was 35 nm. In the case of two-color recording, the red-shifted channel was imaged in a sequential manner on a second camera with a 620/70 nm interval. A 350 W/cm^2^ of 590 nm light per 2 ms was used for the recording. We applied bleaching correction between frames to compensate for switching fatigue. The images presented were deconvolved with a narrow Gaussian of 50 nm FWHM combined with a wider Gaussian of 175 nm FWHM accounting for 10% of the PSF amplitude, such a geometry considers the properties of rsFPs, where a background signal due to a non-photo-switchable fraction of the molecules is expected. The final image is the result of 5 iterations of the Richardson-Lucy algorithm.

### MoNaLISA image analysis

To detect the clusters of Syn1 from the MoNALISA images the following pipeline has been used. First, to identify the clusters, the deconvolved (5 iterations) and bleached corrected images were filtered using a combination of Laplacian filter (3 pixels), to enhance their contrast, and the Gaussian filter (1 pixel), to reduce the background. After a global threshold (common to the whole dataset) the clusters can be defined and parameters like area, intensity and ellipticity extracted. Clusters closer to 70 nm have been merged. To further understand the inner composition of the clusters, we identified the maxima for each cluster (given a tolerance parameter of 5). In the case of two-color recording, the condition of the inhibitory cluster does not depend on the overlap or proximity (less than 70 nm) of the clusters in the two channels.

### hiPSC differentiation to hypothalamic neurons

Control L2135 iPSC line ^85,86^ was differentiated into hypothalamic neurons according to a protocol described before ^87^ with minor modifications. Briefly, iPSCs were maintained in matrigel coated plates with mTeSR1 (Stem Cell Technologies) and medium changes were performed every two days. For hypothalamic neuron differentiation, cells were plated on matrigel coated plates in N2B27 medium (Neurobasal A:DMEM/F12 (1:1) (Life Technologies), 1x B27 without Vit. A (Life Technologies), 1x N2 (Life Technologies), 1x PenStrep, 1x GlutaMAX (Life Technologies), 0,075% sodium bicarbonate (Life Technologies), 0.5 x MEM-NEAA (Life Technologies), 200 nM ascorbic acid (Sigma)) supplemented with SB431542 (Tocris, 10μM), XAV939 (VWR, 2μM) and LDN-193189 (Sigma, 100nM). Medium was exchanged every 2 days with reducing supplement concentration gradually. To ventralize the progenitors 1µM SAG (Merck) and 1µM Purmorphamine (Sigma) were added between day 2 and day 8. DAPT (Tocris, 5 µM) was added from day 8 to day 14. After 14 days, cells were trypsinized and replated with N2B27 medium on poly-D-lysine/laminin-coated 16-18mm coverslips for terminal differentiation in the presence of BDNF (R&D Systems, 10 ng/ml).

### Syt1 nanobody uptake by human stem cell-derived neurons

80-day old human hypothalamic neurons were used for the uptake experiments. All nanobody incubations were performed in BrainPhys medium (StemCell Technologies) to enhance synaptic activity and the diluted nanobodies were centrifuged in a tabletop centrifuge at maximum speed for 10 min at room temperature. The surface Syt1 pool was blocked by incubating the neurons with 4.85 µM unlabeled Syt1 nanobody for 5 min at 37°C in a humid chamber. After washing 3 times with PBS, neurons were incubated with 5.5 µM pHluorin-labelled Syt1 nanobody in the presence of either 30mM KCl or 1µM TTX (Alomone labs) to stimulate neurons or to silence them, respectively, for a total of 20 min at 37 °C in a humid chamber. Cells were extensively washed with PBS, fixed in 4% PFA for 15 min, permeabilized and blocked with 0.01% saponin/10% normal goat/PBS serum and stained with the primary antibodies in the same blocking solution in a humid chamber. Primary antibodies were anti-GFP (Aves labs GFP-1020, 1:1000), anti-Syt1 (Synaptic Systems 105-011; 1:100) and anti-TUBB3 (Biolegend Poly18020; 1:1000). All secondary antibodies were Alexa Fluor-conjugated (ThermoFisher Scientific; 1:500). Nuclei were counterstained with DAPI (Sigma-Aldrich). Images were acquired with the Nikon A1R confocal microscope through a 60X NA1.2 water immersion lens and signal were quantified with Fiji using the Coloc2 plugin (https://imagej.net/plugins/coloc-2).

### iNeuron culture

iNeurons were obtained as described previously ^42^ from the iPSC line BIHi005-A (generated at the Max Delbrück Center). After day 20 neurons were fed weekly, by replacing 1/10 volume of the old media with fresh growth media containing 2 μM Ara-C. For activity measurements, neurons (6-8 weeks in culture) were stimulated with 200 action potentials, 40 Hz 5 s using a field stimulation chamber RC-47FSLP (Warner Instruments) and imaged at physiological temperature (37 °C) in an imaging buffer ^88^ containing 1.3 mM Ca^2+^ with an epifluorescence microscope Nikon Eclipse Ti equipped with a 40X oil objective. Images were acquired every 2 s. Analysis was performed with SynActJ ^89^.

## Acknowledgements

We are deeply grateful to Dr. Kim Ann Saal, Nicole Hartelt, Christina Patzelt, and Christina Zeising for their outstanding technical assistance. We also thank Anita-Karina Jaehnke, Regina Sommer-Kluß and Christin Wiemuth for their continuous organizational support. We thank the beamline scientists at Diamond Light Source (United Kingdom, proposal MX21625) for their valuable support in structural biology data collection. We are grateful to Geoffrey Masuyer for assistance with crystallization and insightful discussions of the results. We also acknowledge Sabine Beuermann and the AGCT Lab for their excellent technical support. This work was supported by the Swedish Research Council (2022-03681, to P.S.), the Novo Nordisk Foundation (NNF20OC0064789, to P.S.), the Swedish Brain Foundation (to P.S.), the Wellcome Trust (204954/Z/16/Z, to M.A.C.), the Australian Research Council (ARC; Discovery Early Career Researcher Award DE190100565, to M.J.), the Deutsche Forschungsgemeinschaft (DFG, German Research Foundation; SFB1286/A01, to B.H.C.; SFB1286/A09, to N.B.; SFB1286/A05, to S.J.; SFB1826/Z04, to F.O.; FOR5228/HA2686/22-1 to V.H.), the European Research Council (ERC) Advanced Grant (SVNeuroTrans, to R.J.; SYNAPSEBUILD (Agreement No. 884281) to V.H.), and the NeuroNex International Network (BR 1107/15-2, to N.B.). C.C. was supported by FWO-SoE postdoctoral fellowship (Ref: 204990/12ZY321N) and N.K. by an EMBO long-term postdoctoral fellowship. The E.F.F. lab received support from a CZI Collaborative Pairs Pilot Project Award (Cycle 2, Phase 1) and was affiliated as an associated member to the SFB1286.

## Author contributions

Author Contributions: Conceptualization: P.S., F.O., R.J., S.O.R., E.F.F. Data Curation: K.J., M.N., H.M.M., F.O., E.F.F. Formal Analysis: K.J., D.I., N.K., K.W., E.S.H., M.J., E.F.F. Funding Acquisition: M.J., M.A.C., P.V., B.H.C., P.S., N.B., R.J., S.O.R., E.F.F. Investigation: R.G., K.J., P.A.G., F.L., M.N., J.H., A.W., S.B., F.P., M.D., N.K., V.K., C.C., V.N.M., N.L.C., E.S.H., H.L., K.W., D.I., T.P.W., C.S., M.J., M.A.C., F.A.M., E.T.K., I.T., B.H.C., F.O., E.F.F. Methodology: R.G., K.J., J.H., A.W., S.B., F.P., M.D., N.K., C.C., K.W., D.I., T.P.W., C.S., M.J., F.A.M., E.T.K., I.T., B.H.C., F.M., V.H., S.J., P.S., F.O., S.O.R., E.F.F. Resources: M.A.C., F.A.M., P.V., N.K., C.C., V.H., S.J., H.U., P.S., F.O., R.J., S.O.R., E.F.F. Supervision: N.L.C., P.V., I.T., E.T.K., V.H., B.H.C., N.B., F.M., S.J., P.S., F.O., R.J., S.O.R., E.F.F. Visualization: R.G., K.J., A.W., S.B., N.K., F.P., K.W., D.I., B.H.C., M.J., E.F.F. Writing – Original Draft: R.G., R.R., N.K., E.S.H., E.F.F. Writing – Review & Editing: All authors. Project Administration: E.F.F.

