## Supplementary Information for "Nanobinders for Synaptotagmin 1 enable the analysis of synaptic vesicle dynamics in rodent and human models"

### Supplementary Figures

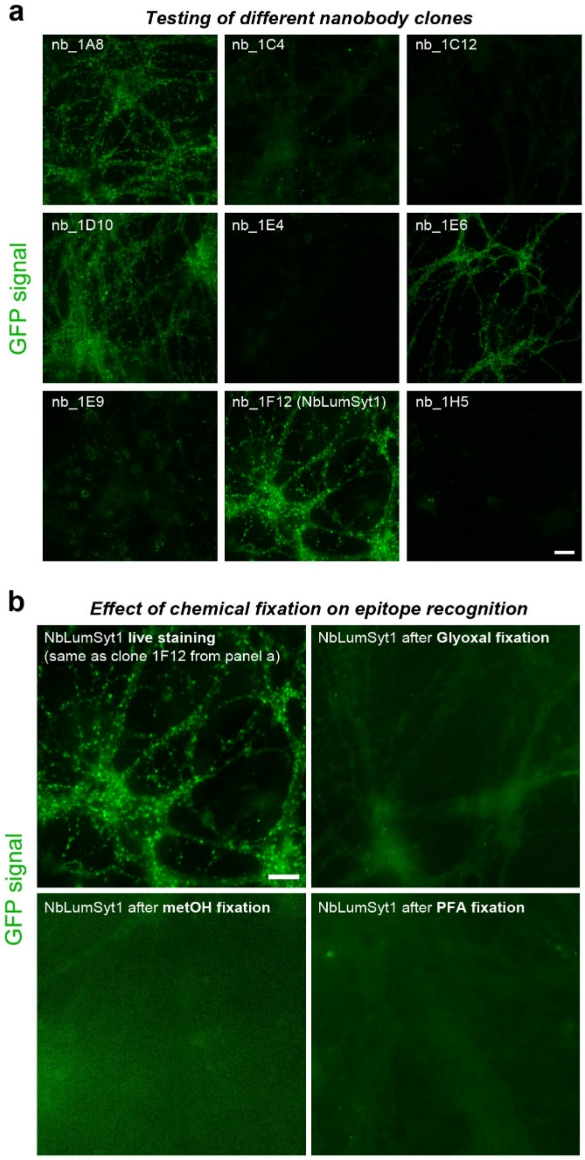


**c** ***Multiple sequence alignement of Syt1 with the epitope bound in light blue***

```
Syt1_DanRe  -- MSRREARVGNPAPTAAPEVPGNSTEAAGPGPRETKDEMFSKVKNKFMNELHKIPLPS    57
Syt1_Human  MVSESHHEALAAPPVTTVATVLPSNATEPASPG--EGKEDAFSKLKEKFMNELHKIPLPP   58
Syt1_Mouse  MVSASRPEALA-APVTTVATLVPHNATEPASPG--EGKEDAFSKLKQKFMNELHKIPLPP   57
Syt1_Rat    MVSASHPEALA-APVTTVATLVPHNATEPASPG--EGKEDAFSKLKQKFMNELHKIPLPP   57

Syt1_DanRe  WAIVAIAFVAVVLVVSCCFCICKKWIFKKKNKKKGKDKG-KNAINMKDVKDGIKT--EAL   114
Syt1_Human  WALIAIAIVAVLLVLTCCFCICKKCLFKKKNKKKGKEKGGKNAINMKDVKDLGKTMKDQA   118
Syt1_Mouse  WALIAIAIVAVLLVVTCCFCVCKKCLFKKKNKKKGKEKGGKNAINMKDVKDLGKTMKDQA   117
Syt1_Rat    WALIAIAIVAVLLVVTCCFCVCKKCLFKKKNKKKGKEKGGKNAINMKDVKDLGKTMKDQA   117
```

***Supplementary Fig. 1: Nanobody clone selection and initial characterizations. a****) Testing the nanobody clones (fused to GFP) that were found to bind the Syt1 epitope by ELISA in live immunofluorescence on primary hippocampal neurons. Among the tested candidates 1A8, 1D10, 1E6 and 1F12 show the most promising binding. After analysis of the protein sequence of the nanobodies these four clones appeared highly related (not shown), hence the clone 1F12, which seemed the strongest binder was selected for further characterizations.* ***b****) Upon chemical fixation the ability to recognize the epitope is lost.* ***c****) Protein sequence conservation analysis of the epitope bound by the nanobody reveals that this region is conserved and presents several lysins (K), which upon fixation due to the reactivity of their primary amine group can form methylene bridges with other surrounding molecules and lose their initial immunogenic properties*

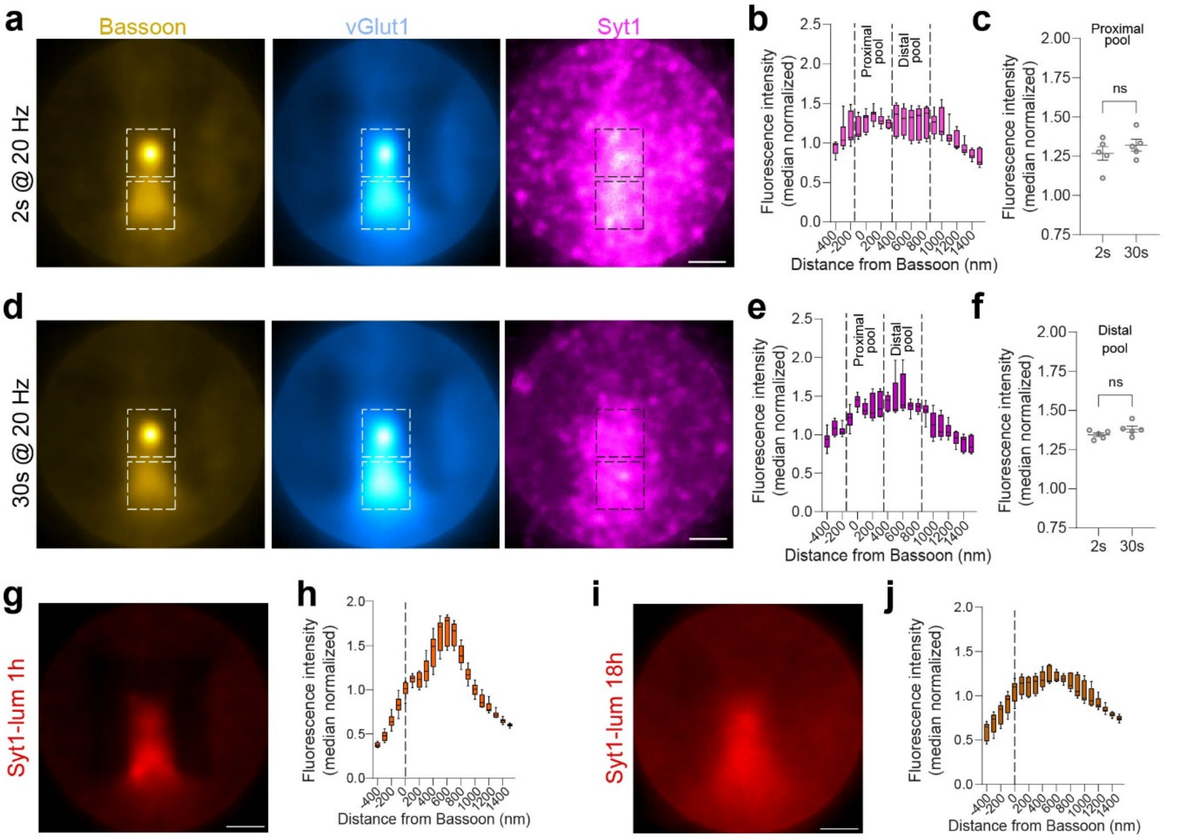


***Supplementary Fig. 2: Distribution of synaptic vesicle populations assessed with a HALO-tag version of the NbLumSyt1.*** *Primary hippocampal neurons were examined in defined experimental modalities to reveal the spatial distribution of SVs following different stimulations and labeling schemes. To exclude effects of Syt1 on the plasma membrane, neurons were first blocked with an unlabeled version of NbLumSyt1 for 5 min at 37ºC. Prior to each manipulation, a fresh aliquot of NbLumSyt1-HALO was pre-mixed with the Janelia Fluor 646 HaloTag ligand for 15 min at RT to ensure efficient labeling.* ***a****) Prior to electrical stimulation, neurons were rapidly washed with a buffer containing 50 µM APV and 10 µM CNQX to avoid recurrent firing and then exposed to the same buffer along with the pre-mixed NbLumSyt1-HALO-JF646. Following the addition of NbLumSyt1-HALO-JF646, neurons were immediately stimulated at 20 Hz for 2 s. In the control condition, neurons were stimulated only in the presence of the unconjugated HaloTag JF646. After stimulation, neurons were allowed to recover for 2 min in the same buffer, washed twice with Tyrode's buffer and fixed. After fixation and quenching, neurons were stained with anti-Bassoon antibody and a vGlut1-STAR580 nanobody. STED images were acquired with constant settings and image analysis was performed using a custom written MATLAB script. Briefly, active zone (Bassoon) spots were detected using preprocessing based on Fourier transformation. The Bassoon spot was considered as the center of the detected presynapses, then the regions were aligned, rotated, and averaged so that the center of the Bassoon signal was placed in the center of the image and the highest intensity was aligned in the lower part of the image (see yellow leftmost panel in a). VGlut1 and NbLumSyt1-HALO-JF646 images were aligned using the same parameters as the Bassoon reference image, and a vertical intensity line profile was defined and intensity measured vertically along the averaged Bassoon peak. The line profile was measured for each coverslip separately and the raw intensity profile values for each coverslip were median normalized, plotted (****b****) and quantified (****c****). This analysis allowed to clarify that the proximal pool of SVs (arbitrarily defined as vesicles closer than 300 nm from the Bassoon center), is similarly distributed for both 2s stimulations and 30s stimulations (panels* ***d*** *and* ***e****). Likewise, the distal pool of SVs (arbitrarily defined as vesicles between 400 and 800 nm from the Bassoon center) show no striking difference between stimulation regimes (****f****). Collectively, these data suggest that there is rapid mixing of SVs between proximal and distal pools during recycling, and that different stimulus trains do not recruit functionally distinct types of SVs. Similarly, to experiments in* ***a-f****, uptake assays were designed to evaluate the distribution of "younger" (1h-old;* ***g-h****) and "older" (18h-old;* ***i-j****) SVs. After 30 min labeling of primary hippocampal neurons with NbLumSyt1-HALO-JF646, the neurons were followed for different times and images were taken and analyzed as in* ***a-f****. These analyses show a broadening of the NbLumSyt1-HALO-JF646 peak, which likely reflects less efficient localization of aged SVs and possibly the relocalization of aged Syt1 molecules to other cellular compartments. Scale bars: 500 nm*

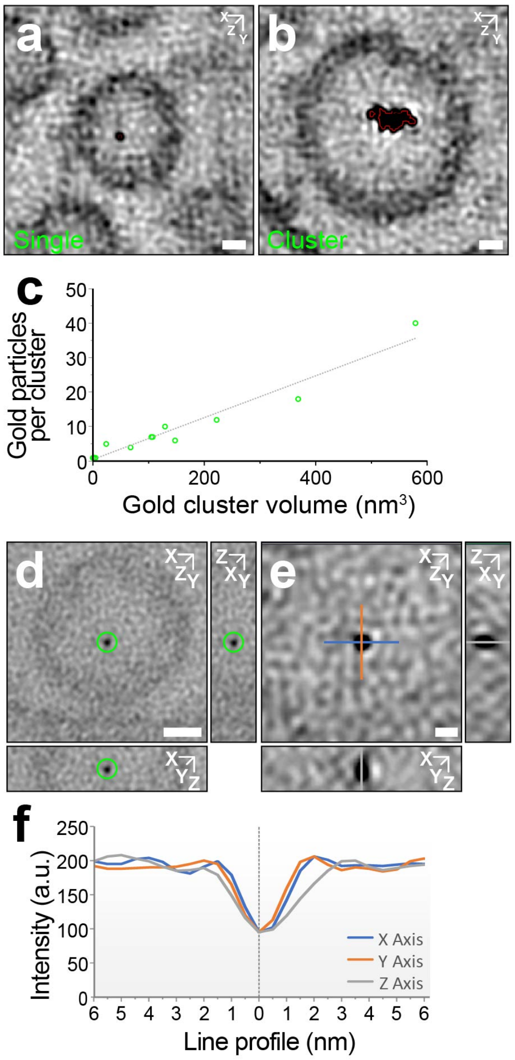


***Supplementary Fig. 3: Additional measures relating to main Figure 2. a, b**) Electron tomographic sub-volumes acquired at 36 000 x magnification (binned voxel size x,y,z = 1.2 nm; projection through 5 tomographic slices) illustrating gold-labelled clear-core synaptic vesicles (CCSV$^{G}$) from the NbLumSyt1-ALFA-tag:NbALFA-Gold complex (NbLumSyt1) treated synapses. A single luminal gold particle (a) and a cluster (b) are shown (red outline indicates automatic segmentation of gold used to estimate luminal gold-label volumes. **c**) Plot demonstrating the linear relationship between the number of manually quantified luminal gold particles and corresponding unbiased volume estimates in individual vesicles from a tomogram acquired from an NbLumSyt1-ALFA-tag:NbALFA-Gold complex treated synapse. **d-f**) Electron tomographic sub-volume acquired at 57 000 x magnification (unbinned voxel size x,y,z = 0.5 nm; projection through 6 tomographic slices) from an NbLumSyt1-ALFA-tag:NbALFA-Gold complex treated synapse in which a single luminal gold particle is clearly resolved (**d**). An enlargement (**e**) indicates orthogonally positioned line profiles centered on the gold particle (blue, x-axis; orange, y-axis; grey, z-axis). Corresponding intensity profiles (**f**) consistent with the detection of an individual 3 nm gold particle. Scale bars: a, b, d 10 nm; e, 3 nm.*

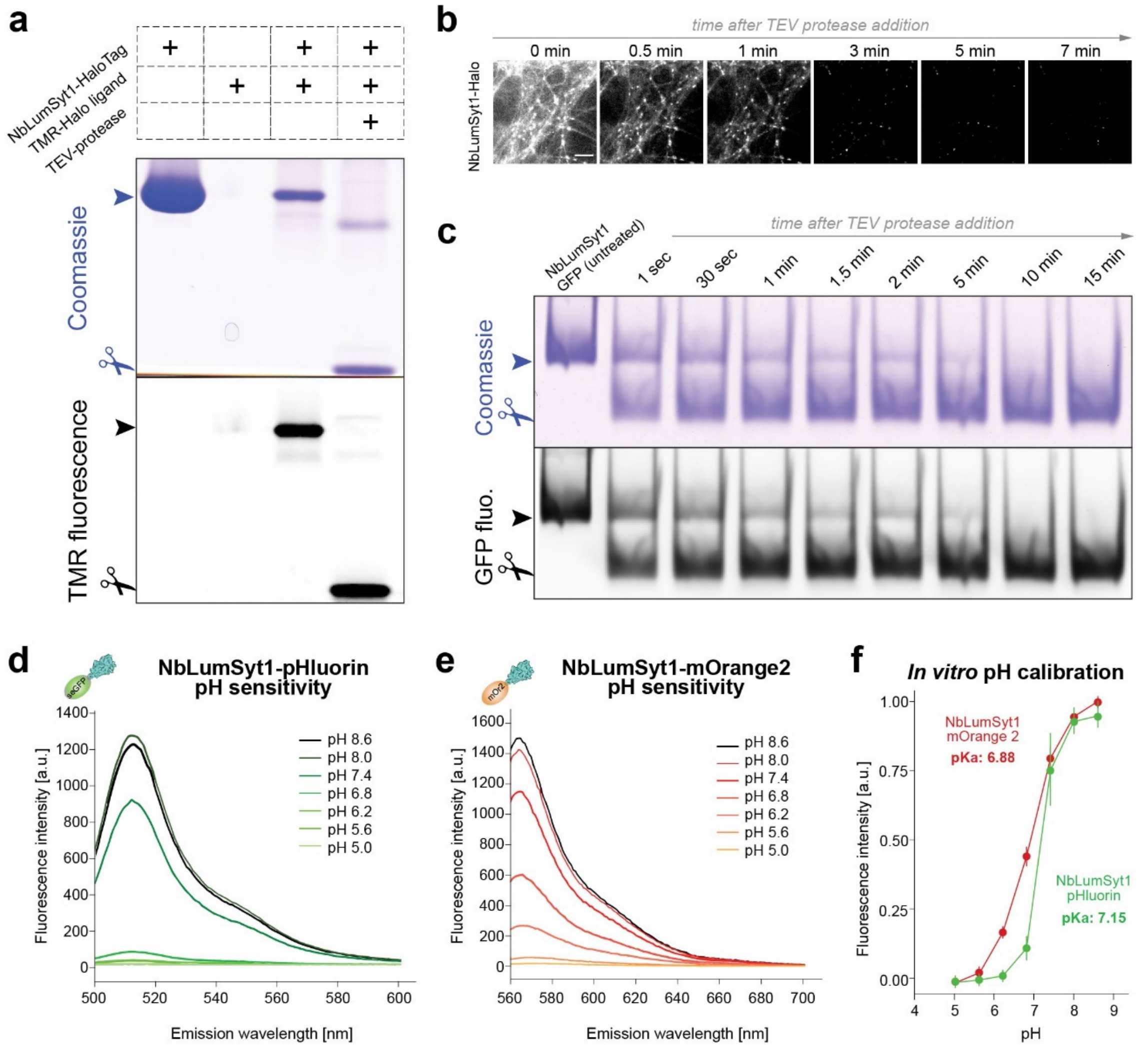


***Supplementary Fig. 4: TEV protease-cleavable versions of NbLumSyt1 for studying the readily-retrievable pool of synaptic vesicles.*** ***a****) Analysis by SDS-PAGE of the NbLumSyt1-TEV-Halo, either alone or in complex with TMR-labeled Halo ligand, before and after TEV protease treatment. Upon TEV cleavage the NbLumSyt1-TEV-Halo/TMR ligand complex reveals two bands, with the lower one containing the Halo-Tag/Halo ligand complex and is fluorescent (lower image), and an upper band containing the separated nanobody.* ***b****) Upon live labelling of primary neurons with NbLumSyt1-TEV-mOrange (surface only), and TEV-protease treatment, the mOrange signal is efficiently removed from cultured neurons, showing that the fluorescent signal is derived from surface exposed pools since the intravesicular pools are quenched due to their low pH.* ***c****) Similar experiment as in a), performed in isolated NbLumSyt1-TEV-mOrange reveals that the time course of cleavage resembles that of labeled neurons.* ***d-f****) Estimation of resting pH using the NbLumSyt1-pHluorin (****d****) or the NbLumSyt1-mOrange2 (****e****) in isolated molecules (in vitro). For the same measurement in primary neurons refer to the main figure. To estimate the pKa values, calibration curves were created by titrating the pH-response of the probes. In accordance with previous reports [28], the pKa of mOrange is lower than that of pHluorin, extending the measuring range towards lower pH-values.*

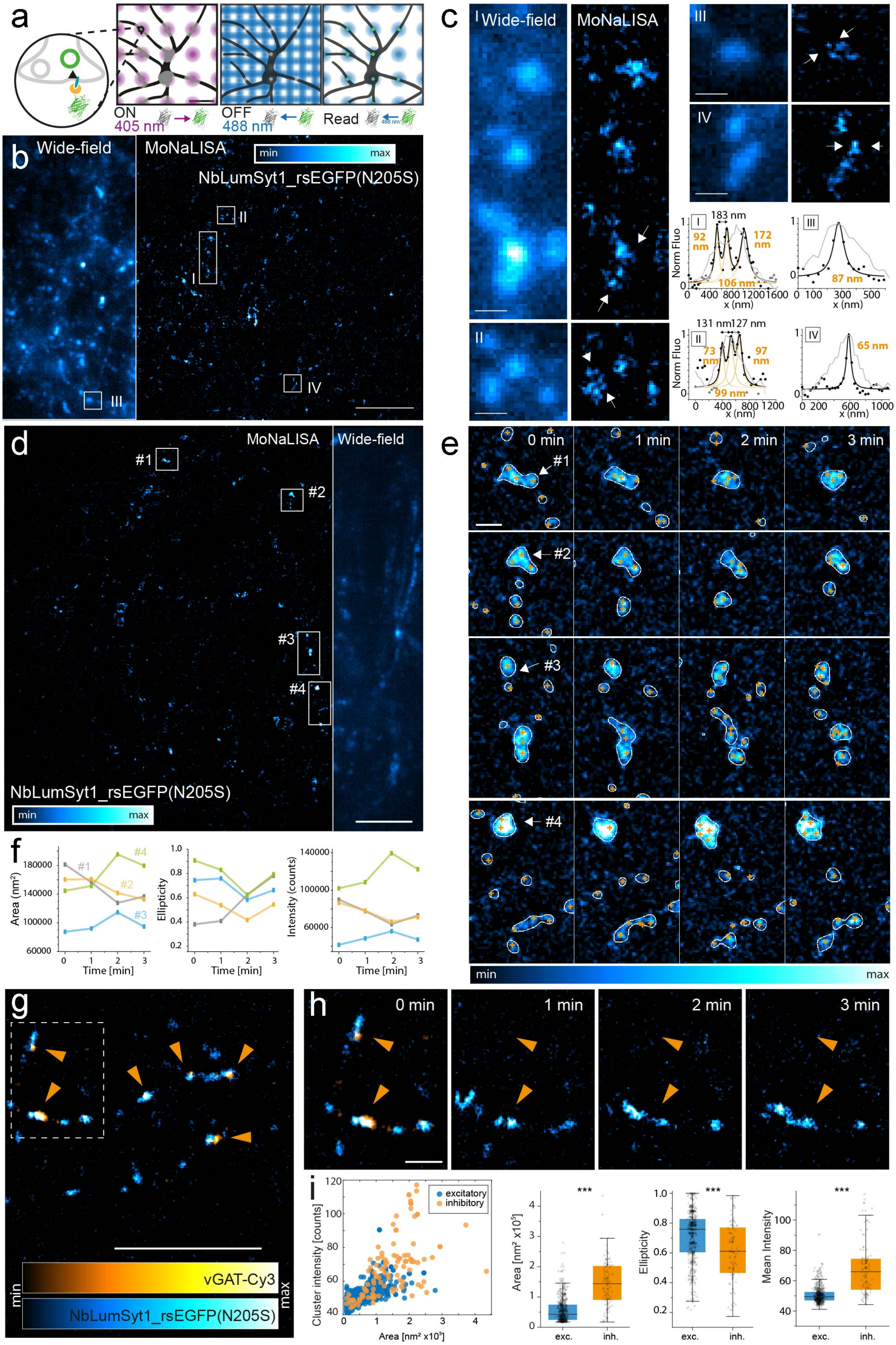


***Supplementary Fig. 5: MoNaLISA diffraction unlimited imaging of NbLumSyt1-rsEGFP(N205S). a)*** *Schematic illustration of the MoNaLISA imaging scheme. The illumination is parallelized over an extended field of*

*(continuing from previous page)*

*view using multi-foci to switch in the ON-state the rsFPs (reversibly switchable Fluorescent Proteins) located in an array of focal spots and sinusoidal light patters to switch the rsFPs located in the periphery of each of the foci in the OFF state. The fine regions containing the remaining rsFPs in the ON state are then read-out with an additional array of blue foci.* ***b, c****) Comparison between a wide-field image and a MoNaLISA image for live hippocampal neuron labelled with NbLumSyt1-rsEGFP(N205S). The ROIs in c show different zoom-in around different regions of the field of view, reporting the resolution achieved in the visualization of the cluster through lines profile traced across the white arrows. Clusters of sizes down to 65 nm in width can be isolated with separation down to 130 nm.* ***d, e****) Time lapse imaging of the NbLumSyt1-rsEGFP(N205S) at intervals of 1 min between the images.* ***f****) Syt1 clusters can increase (#4) or decrease (#1) in area, number and overall shape of the active zone within minutes. The graphs follow area, ellipticity and intensity for 4 clusters out of the 73 identified clusters in the same field of view.* ***g-i****) Dual color imaging and quantifications with NbLumSyt1-rsEGFP(N205S) in the cyan channel and an inhibitory synapses marker in the orange channel (vGAT-Cy3).* ***i****) Inhibitory synapses (vGAT-positive), show a significant increase in area and mean intensity. The shape of the active zone of inhibitory synapses also resulted mostly elliptical and packed with an increased number of small clusters. Ellipticity is defined as the ratio between the minor and the major axis of each cluster (ranging from 0 to 1). The graphs are the result of 648 clusters on 6 experiments, with a mean of around 100 clusters identified per field of view. Unpaired Student's t-test. *** = p ≤ 0.001. Scale bars: a 650 nm; b, d, g 5 µm; c, e 500 nm; h 1 µm.*
